# Disruption to rRNA biogenesis triggers histone mRNA 3’ misprocessing in *C. elegans* through genome-surveillance regulators

**DOI:** 10.64898/2026.11.16.766232

**Authors:** Mahsa Golshani Nasab, Spencer R. Dmytruk, Anna C. McCullough, Cheng-Wei Wu

**Affiliations:** Department of Veterinary Biomedical Sciences, Western College of Veterinary Medicine, University of Saskatchewan, Saskatoon, SK, S7N 5B4, Canada; Toxicology Centre, University of Saskatchewan, Saskatoon, SK, S7N 5B3, Canada; Department of Biochemistry, Microbiology and Immunology, College of Medicine, University of Saskatchewan, Saskatoon, SK, S7N 5E5, Canada

**Author notes:** For correspondence (CWW).

## Abstract

Maturation of replication-dependent histone pre-mRNA transcripts involves a 3’ cleavage event facilitated by the conserved stem-loop binding protein (SLBP). Cellular conditions that trigger an error in this process are not well understood. Here, we developed a GFP-based *in vivo* histone 3’ misprocessing reporter in *Caenorhabditis elegans* and performed a genome-wide RNAi screen for histone regulators. We found that depletion of genes involved in various steps of ribosomal RNA (rRNA) biosynthesis triggers misprocessing of histone 3’ tail, with the strongest effect observed upon depletion of *rpoa-2* encoding the core subunit of RNA polymerase I. Transcriptome sequencing reveals *rpoa-2* knockdown mirrors the effect of depleting the *C. elegans* SLBP homologue CDL-1 at the protein level, indicating that *rpoa-2* and CDL-1 co-regulate similar transcriptome targets. Depletion of *rpoa-2* impairs endomitosis of the *C. elegans* intestinal cells and is accompanied by aberrant activation of cell cycle markers in the absence of change in DNA content. Gene interaction analysis shows that *rpoa-2* signals through multiple conserved cell cycle regulators, including ATM-1, GEN-1, and LIN-35 to trigger histone misprocessing in both CDL-1 dependent and independent manners. Together, our findings reveal that disruption in the rRNA biogenesis pathway triggers histone misprocessing through conserved genome surveillance pathway regulators.

**Graphical abstract:** 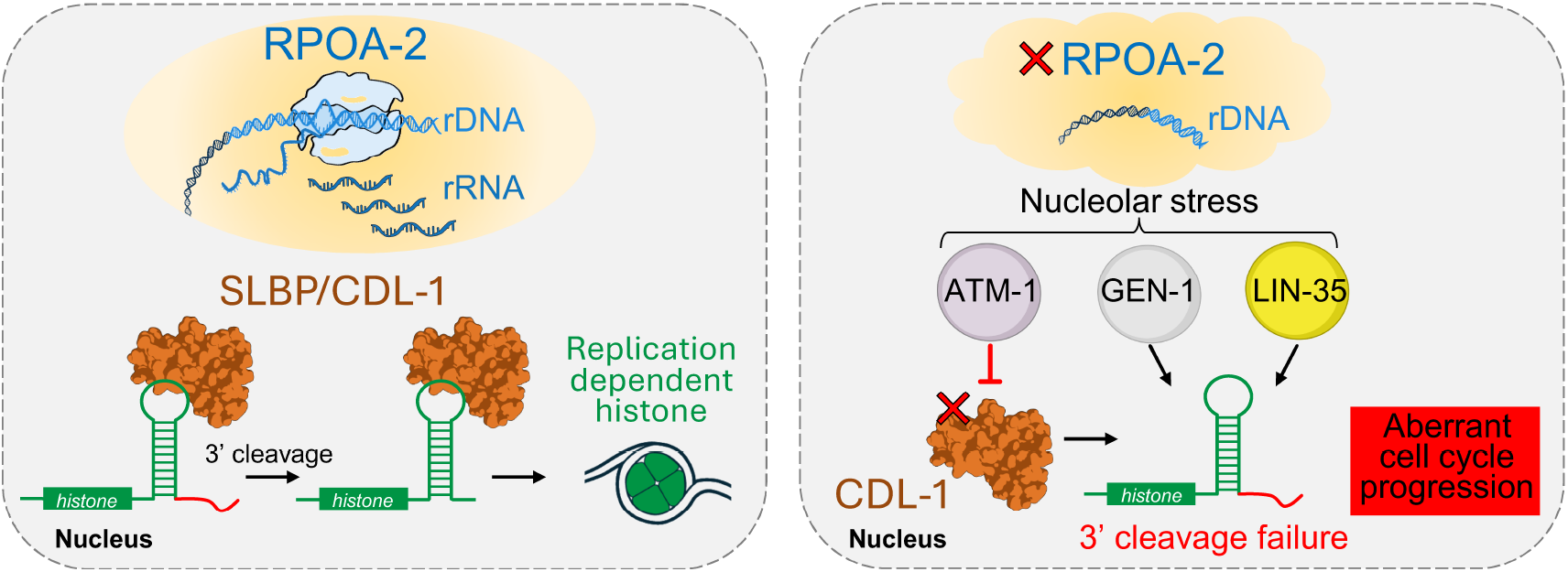

## Introduction

Histone proteins are found in all eukaryotic organisms as they form the fundamental components of chromatin that package DNA. Replication-dependent histone proteins are produced in synchrony with DNA replication to create 4 core variants (H2A, H2B, H3, and H4) to make up the nucleosome (1). A unique feature of replication-dependent histones is that they are intronless and are the only mRNA that is not polyadenylated after transcription; rather, the histone 3’ UTR forms a conserved stem loop that recruits various cleavage factors that drive transcript maturation via 3’ cleavage (2, 3). This cleavage process is akin to the evolutionarily conserved post-transcriptional 3’ cleavage of snRNA transcripts required for their maturation, which is catalyzed by the Integrator complex (4–6). In mammals, the principal protein that initiates histone 3’ cleavage is SLBP (stem loop binding protein) that recognizes and interacts with the conserved stem loop located 15-50 bp downstream of the coding region (7). This is followed by binding of the U7 small nuclear RNA (snRNA) to the adjacent histone downstream element (HDE) motif to recruit various cleavage factors to catalyze endonucleolytic processing of the histone tail (3). Upon successful 3’ cleavage, the mature histone mRNA remains bound to SLBP and is exported to the cytoplasm for protein synthesis followed nuclear re-entry (3). While the SLBP-U7 dependent mechanism of histone processing is conserved in metazoans, the nematode *Caenorhabditis elegans* is an exception. *C. elegans* expresses the conserved SLBP homologue called CDL-1 (cell death lethal) that binds to the 3’ stem loop to facilitate histone 3’ processing, but lacks the canonical U7 sRNA gene for HDE recognition (8, 9). An alternative mechanism has been proposed in *C. elegans* where an Argonaute encoding protein called CSR-1 (Chromsome-Segregation and RNAi deficient) promotes 3’ processing via binding to siRNAs that are complementary to sequences downstream of the histone stem loop to guide cleavage (10, 11). While CSR-1 exhibits endonuclease activity, whether it directly cleaves the histone 3’ tail has yet to be demonstrated.

Despite mechanistic differences in U7 requirement, SLBP has been shown to be the central regulator of histone processing. Disruption of SLBP results in histone 3’ misprocessing that is characterized by transcriptional read-through of the histone 3’ UTR, resulting in aberrant polyadenylation driven by the usage of cryptic polyadenylation signals located downstream of the HDE. (9, 12). While histone mRNA generated by read-through errors is stabilized by polyadenylation, histone proteins are generally down-regulated when SLBP is disrupted (10, 13). This is likely due to the continuous requirement of SLBP in histone protein production from mRNA nuclear export to interaction with the translational complex (1). Consistent with the essential role of histones, loss of SLBP is lethal across *C. elegans, Drosophila*, zebrafish, and mouse (8, 14–16). At the cellular level, loss of SLBP dependent histone production is associated with genomic instability, impaired DNA replication, and chromosome condensation failure (14–16). More recently, mutations that impair proteins associated with the histone processing machinery have been linked to human pathologies including autoinflammatory disorders, spinal muscular atrophy, and neurodegenerative diseases (17–19).

To date, while core components of the histone cleavage complex are well described (3), with the exception of *C. elegans*, genetic perturbations or sensing mechanisms that trigger histone misprocessing are not well understood. In the context of therapeutic discovery, identifying genes that are required for histone processing could reveal new targets to impair S-phase progression of rapidly dividing cells, by limiting the availability of histones required for supporting nucleosome assembly after DNA replication (20).

Here, we adapted our previous construction of a snRNA misprocessing reporter to develop a GFP based *in vivo* histone 3’ misprocessing reporter in *C. elegans* and performed a genome-wide RNAi screen to search for gene silencing conditions that impair histone pre-mRNA processing (5). We identified a novel role for the ribosomal RNA biogenesis pathway in maintaining histone pre-mRNA processing, with disruption to *rpoa-2* (RNA Pol I A subunit) encoding the largest subunit of RNA polymerase I causing widespread histone 3’ misprocessing. The *C. elegans* RPOA-2 is responsible for the transcription of the polycistronic 47s rRNA precursor that is processed into 18S, 5.8S, and 28s rRNA which is assembled into the ribosome to form an active translational complex (21). While *rpoa-2* is best characterized for its requirement in development and longevity in *C. elegans*, recent studies have also implicated this gene in contributing to chromatin accessibility and germ cell apoptosis (22–25). Here, we show that *rpoa-2* co-regulates a substantial share of CDL-1 dependent gene expression, and that depletion of *rpoa-2* triggers a defect in intestinal endomitosis that is accompanied by aberrant expression of cell cycle markers. Mechanistically, we show that *rpoa-2* disruption signals through multiple conserved downstream effectors to induce histone 3’ misprocessing, including ATM-1 (Ataxia-telangiectasia mutated homologue), GEN-1 (GEN1 Holliday junction resolvase homologue) and LIN-35 (Retinoblastoma homologue). This indicates that rRNA biogenesis disruption signals through the DNA damage response pathway and negative cell cycle regulators to impair histone pre-mRNA processing. Taken together, this study provides new insights into a crosstalk between rRNA biogenesis and histone post-transcriptional regulation through genome-surveillance factors.

## Materials and Methods

### C. elegans strains and construction

All *C. elegans* strains were maintained using standard methods at 20°C, unless mentioned otherwise (26). The following strains where used: N2 Bristol wildtype, MWU235 *cwwIs14[his-6p:his-6::GFP; myo-2p::tdTomato],* PHX10868 *syb10868*[*cdl-1::AID*::3xFLAG:mNeonGreen]*, MWU258 *syb10868*; *ieSi38 [sun-1p::TIR1::mRuby::sun-1 3’UTR + Cbr-unc-119(+)]; wrdSi23 [eft-3p::TIR1::F2A::mTagBFP2::AID*::NLS::tbb-2 3’UTR] (I:-5.32),* MWU259 *ieSi38 [sun-1p::TIR1::mRuby::sun-1 3’UTR + Cbr-unc-119(+)]; wrdSi23 [eft-3p::TIR1::F2A::mTagBFP2::AID*::NLS::tbb-2 3’UTR] (I:-5.32)*, ESC332 *cse319[AID*::GFP::rpoa-2]* I; *ieSi57 [eft-3p::TIR1::mRuby::unc-54 3’UTR + Cbr-unc-119(+)] II,* DQB298 *bmdSi86 [LoxN::rps-27p::DHB::GFP::P2A::H2B::mKate2] I*, TJ1 *cep-1(gk138) I*, GOU3941 *his-48(cas848[gfp::his-48 knock-in]) IV*, VC381 *atm-1(gk186) I*, TG1540 *gen-1(tm2940) III,* AWR58 *lin-35(kea7[lin-35p::AID*::GFP::lin-35]) I; keaSi10 II*.

The MWU235 *his-6* misprocessing strain was developed by adapting a cell based reporter systems previously described in *Drosophila* and an *in vivo* read-through reporter we previously developed in *C. elegans* (5, 12). In brief, we first synthesized an 835 bp DNA fragment via Genestrand (Eurofins Genomics, Louisville, KY) comprising of the *his-6* transcript promoter, *his-6* coding region with stop codon removed, *his-6* 3’ downstream sequence, followed by the 51 base pairs of the multiple cloning site sequence of the pPD95.75 GFP plasmid. Fusion PCR was then used to combine the synthesized *his-6* fragment in frame with the GFP sequence containing the *unc-54* 3’UTR from the pPD95.75 plasmid (27). The construct was microinjected into *C. elegans* together with the co-injection *myo-2*::tdTomato marker. The expressed extrachromosomal array was then stably integrated into the genome via U.V. irradiation. The integrated strain was outcrossed 6 times with wildtype N2 before use. The CRISPR generated insertion of AID*::3xFLAG:mNeonGreen into *cdl-1* was obtained from Suny Biotech (Fuzhou, China), with single worm PCR used to genotype the insertion using primers listed in **Table S1**. Insertion sites were confirmed with Sanger sequencing.

### Genome-wide RNAi screen and RNAi experiments

An unbiased genome-wide RNAi was carried out using protocols previously described in detail with slight modifications (28). In brief, synchronized L1-stage MWU235 larvae were obtained by hypochlorite treatment and grown in liquid nematode growth media (NGM) and fed with HT113(DE3) bacteria producing dsRNA for 48 hours. After 48 hours of incubation, residual bacteria were removed with a MultiFlo FX plate washer fitted with an AMS module, with worms resuspended in M9 buffer containing 10 mM levamisole. Paralyzed worms were then imaged using a Cytation-5 system to detect GFP fluorescence as an indicator of histone transcript misprocessing. A total of ∼19,000 dsRNA clones were screened and obtained from the MRC genomics RNAi feeding library (Geneservice, Cambridge, UK) and the ORFeome RNAi feeding library (Open Biosystems, Huntsville, AL). Genes showing strong GFP signals were validated twice in a secondary screen on solid NGM agar plates for confirmation. For solid RNAi experiments, NGM agar plates contained 50 µg ml ^-1^ carbenicillin and 100 µg ml ^-1^ of isopropyl β-D-thiogalactopyranoside (IPTG) and were seeded with HT115(DE3) *E. coli* expressing the corresponding dsRNA. Bacteria on NGM agar plates were allowed to grow for 2 days at room temperature prior to use. For all RNAi experiments, the pPD129.36(L4440) plasmid was used as the empty vector (EV) control.

For creating RNAi clones to individually knock down 5.8s, 18s, and 26s rRNA, Q5 High-Fidelity Taq polymerase (NEB, M0492S) was used to amplify DNA fragments of *rrn-2.1* (112 bp), *rrn-1.1* (874 bp), and *rrn-3.1* (1288bp) using primers with *attB1* and *attB2* overhangs, followed by Gateway cloning into the pGC31 RNAi feeding plasmid (29). Primers used for RNAi cloning are listed in **Table S1**. RNAi clones were verified using Sanger sequencing.

### Microscopy and fluorescent analysis

Microscopic analysis was performed using a Zeiss Axioskop 50 fitted with a Retiga R3 camera and Zeiss AttoArc HBO 100W lamp illuminator. Worms were mounted onto a glass slide containing 2% agar pad and immobilized with 2% sodium azide dissolved in M9 buffer. For all experiments, worms were synchronized and transferred at L1-stage onto the desired RNAi or regular NGM plates and imaged after 48 hours, corresponding to development in L4/young adults. Images were taken in greyscale and colourized in ImageJ; the merge channel function was used to create composite images expressing multiple fluorophores. Quantification of fluorescence intensity was performed in ImageJ using the measure function, with corrected total cell fluorescence (CTCF) calculated as integrated density – (area of selection x mean background fluorescence). Background fluorescence was determined for each image by measuring an area on the micrograph where fluorescence was absent.

For total PCN-1::GFP and CDL-1::mNG quantification, total CTFC was calculated using the fluorescent signal for the midbody region as fluorescence was not expressed in the anterior/head and posterior/tail part of the worm. For PCN-1::GFP nuclear/total ratio, CTFC of intestinal nuclei was measured separately with the ratio expressed as nuclei CTFC / (total CTFC – nuclei CTFC). DHB::mKate2 quantification the ratio expressed as cytoplasmic (total CTFC – nuclear CTFC) / nuclear CTFC. For intestinal DNA fluorescence, CTFC was calculated in the anterior intestinal nuclei. Posterior intestinal nuclei were not quantified as images were focused on the anterior intestine, causing the posterior nuclei to fall out of the focal plane. All experiments were performed in 2 independent trials, with the number of animals quantified for each dataset indicated in the figure legends.

### RNA extraction, qPCR, mRNA-sequencing

For each experimental condition, total RNA was extracted from four biological replicates using the PureLink RNA Mini Kit (ThermoFisher, 12183025). Each replicate consisted of approximately 800–1,000 worms, which were lysed using a Qsonica Q55 sonicator. For cDNA synthesis, isolated RNA samples were first treated with DNase I (ThermoFisher, EN0521) to remove residual genomic DNA. Using the Invitrogen MultiScribe reverse transcriptase system (ThermoFisher, 4311235), cDNA was synthesized with random primers on an Applied Biosystems ProFlex thermal cycler following the manufacturer’s protocol. Quantitative PCR (qPCR) was performed on a QuantStudio 3 Real-Time PCR system using PowerUp SYBR Green Master Mix (ThermoFisher, A25741). Relative transcript abundance was calculated after normalization to the reference housekeeping gene *cdc-42*. We validated *cdc-42* as a viable housekeeping gene in that depletion of *rpoa-2* by RNAi, CDL-1 by auxin, or the combination of both, had no effect on *cdc-42* expression relative to the control condition (**Fig. S1a**). Compared to the panel of total and misprocessed histone transcripts measured, *cdc-42* was the most stable gene as determined by BestKeeper analysis, indicating that it is a stable housekeeping gene even under perturbations that cause cell cycle disruption (30) (**Fig. S1b**). Primers used for PCR are listed in **Table S1**.

**Figure S1.**
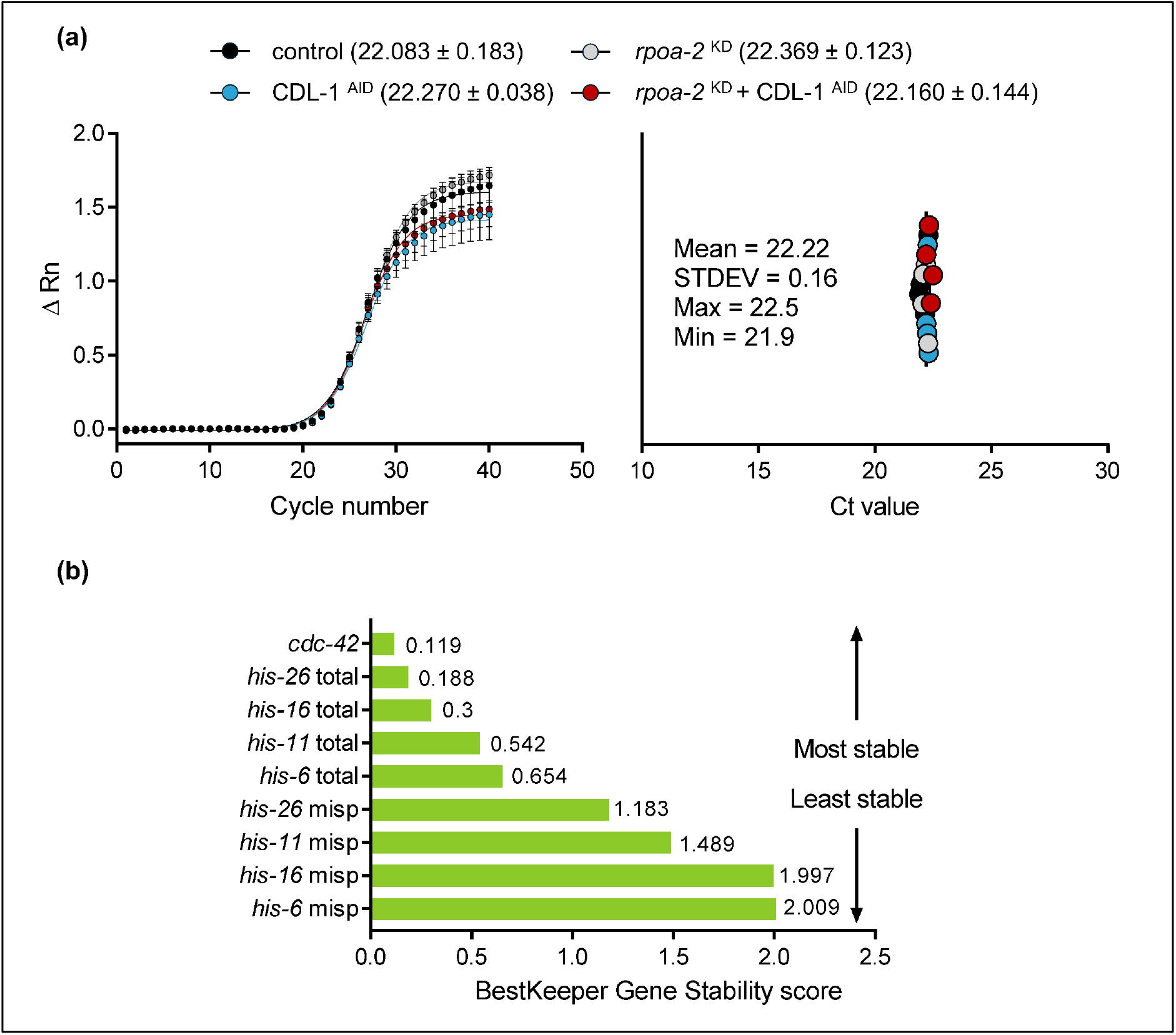
*cdc-42* expression is stably expressed. **a)** qPCR amplification curve and Ct value of *cdc-42* in control, *rpoa-2* ^KD^, CDL-1 ^AID^, or *rpoa-2* ^KD^ + CDL-1 ^AID^ double knockdown RNA samples. **b)** Bestkeeper gene stability analysis of *cdc-42* and total or misprocessed histone variants (30).

For mRNA sequencing, RNA was extracted using the same methods described above, with 3 biological replicates prepared and sequenced for each condition. Frozen RNA samples were delivered on dry ice to GIFS Ag-Tech (Saskatchewan, SK) to perform library construction with oligo(dT) enrichment followed by sequencing on the Element BioScience AVITI system. Sequence reads were mapped to the Wbcel235 genome with STAR, and gene quantifications were performed with RSEM, with DESeq2 used to analyze differential expression (31–33). PCA analysis was performed with ClustVis and Gene Ontology analysis was performed with DAVID functional enrichment (34, 35). The MWU259 strain expressing only the TIR1 proteins was also sequenced in parallel to determine non-specific genes that respond to auxin treatment, which were removed from the MWU258 auxin analysis. Organized data from RNA-seq are shown in **Table S2**.

### Whole worm DNA staining

Synchronized L1 wildtype N2 worms were fed with EV or *rpoa-2* RNAi until they reached the L4 stage and prepared for DNA staining with Hoechst 33342 (ThermoFisher, H3570) following protocols described previously (36). Briefly, worms were washed with M9 to remove residual bacteria and permeabilized with PBST + 0.1% Triton X-100 followed by fixation with Carnoy’s solution overnight at 4°C. The next day, worms were rehydrated with 70% ethanol and washed with PBS followed by incubation with 500ng/ul Hoechst 33342 (ThermoFisher, H3570) for 1 hour at 37 °C. Worms were then washed 3 times with PBST and mounted for fluorescent imaging with the DAPI filter using steps described above.

### Actinomycin D treatment

A 20 mg/mL stock solution of Actinomycin D (ThermoFisher, A7592) was prepared in DMSO (Fisher Scientific, BP2311) and diluted to 100, 150, 200 µg/mL with 5x concentrated OP50 bacteria. An aliquot of 1 mL of each concentrated OP50 mixed with Actinomycin D was seeded on a 6 cm NGM plate and left to dry overnight prior to use. OP50 mixed with 1% DMSO served as the control. Synchronized MWU235 L1 worms were grown on the seeded plates for 48 hours prior to fluorescent imaging. For qPCR experiments, synchronized N2 worms were grown on plates seeded with OP50 mixed with 200 µg/mL Actinomycin D or 1% DMSO for 48 hours, followed by RNA extraction and cDNA synthesis using steps described above.

### Statistical analyses

Statistical analyses were performed in GraphPad Prism 7.05 or 8.4.3 (La Jolla, CA). Student’s t-test with two-tailed P-value calculation was used for comparing two groups, one-way ANOVA with Holm-Sidak post hoc test was used for comparing more than two groups, two-way ANOVA with Holm-Sidak post hoc test was used for two factor comparisons. Correlation was analyzed using the Pearson’s correlation test. Statistical significance is denoted by *P<0.05, **P<0.01, and ***P<0.001. All P values for RNA-seq analyses were corrected for FDR.

## Results

### Construction of an in vivo histone misprocessing reporter

The *C. elegans* histone mRNA expresses the 3’ stem loop structure as conserved in other metazoan species, however, no consensus histone downstream element (HDE) motif has been identified. To generate an *in vivo* histone misprocessing reporter as a biomarker for cleavage error, we first aligned the 3’ nucleotide sequence downstream of the stem loop for all replication-dependent histone transcripts. Alignment of sixty-one replication-dependent histones encoding the four core variants revealed a consensus motif of AAATCC, positioned 13 to 18 nucleotides downstream of the stem loop and upstream of a putative poly-A signal (**Fig. 1a**). This consensus motif has previously been proposed as a putative HDE (9). Alignment of the downstream sequence for individual histone variants shows that this motif is highly conserved for H2A, H2B, and H4, with higher variability observed for H3 (**Fig. S2**). In mammalian histone transcripts, 3’ cleavage occurs between the stem loop and the HDE, as such, we used the position of the AAATCC motif as a reference to approximate the site of histone cleavage in *C. elegans* (3). We synthesized a DNA fragment encoding *his-6* promoter, *his-6* coding region with the stop codon removed, and the 3’ non-coding region with GFP fused nine nucleotides downstream of the putative HDE motif prior to the poly-A signal. While the putative HDE for *his-6* was relatively divergent from the consensus sequence, it was chosen as a candidate gene to construct the *in vivo* misprocessing reporter as the 3’ non-coding region does not encode any stop codon sequence, which is required for GFP to be expressed in frame with the HIS-6 protein to create a fusion construct to detect read-through errors (**Fig. 1b**). The construct was introduced into *C. elegans* via microinjection into the germline followed by UV irradiation to integrate the array into the genome for stable expression.

**Figure 1.**
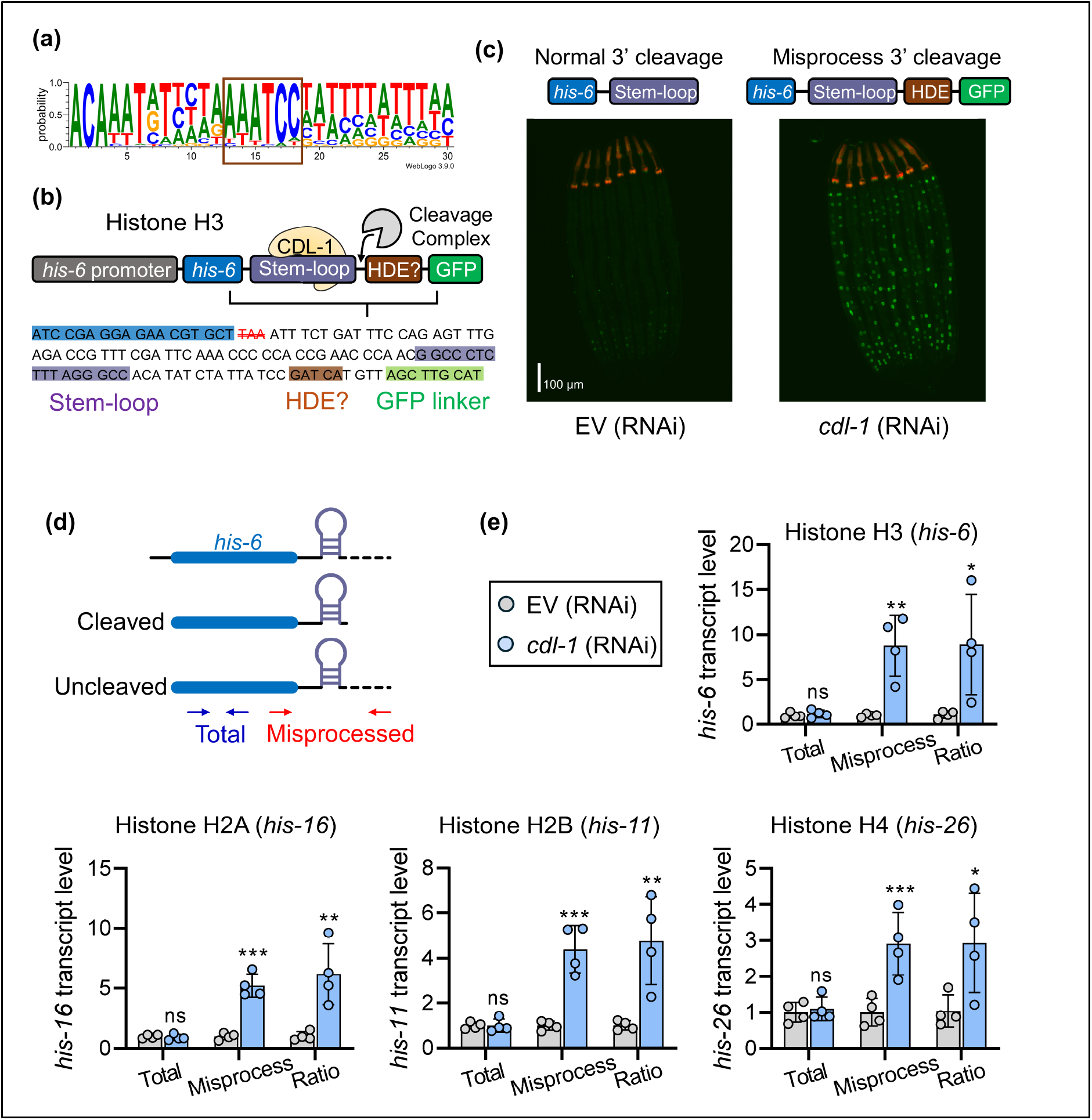
Construction and validation of an *in vivo* histone mRNA 3’ misprocessing reporter. **a)** Sequence logo of nucleotides downstream of the stem loop for sixty-one replication-dependent *C. elegans* histones. Numbers on the x-axis indicate nucleotide position downstream of the stem loop. A consensus motif is highlighted as a potential *C. elegans* HDE. **b)** Schematic of *his-6* 3’ misprocessing reporter. Synthesized fragment of *his-6* promoter, transcript, and 3’ sequenced was fused in frame with GFP and integrated into the *C. elegans* genome to create an *in vivo* histone misprocessing reporter. Stop codon was removed as indicated by the red double strikethrough font. Nucleotide positions of the 3’ stem loop, putative HDE, and GFP linker are shown. **c)** Fluorescent micrograph illustrating activation of the histone misprocessing reporter after RNAi depletion of *cdl-1* that serves as the principal binding protein to the 3’ stem loop. The orange signal near the pharynx represents the *myo-2*::tdTomato co-injection marker. **d)** Primer design to distinguish total and misprocessed histone transcripts. The reverse misprocessed primer binds to regions downstream of the predicted cleavage site between the stem loop and HDE. **e)** Relative levels of total, misprocessed, and misprocessed/total ratio of transcripts encoding histone H3 (*his-6*), histone H2A (*his-16*), histone H2B (*his-11*), and histone H3 (*his-26*). Bar graphs show mean ± STDEV with N = 4 biological replicates measured for each condition; ns: not significant, *P<0.05, **P<0.01, ***P<0.001 as determined by student’s t-test relative the EV(RNAi).

**Figure S2.**
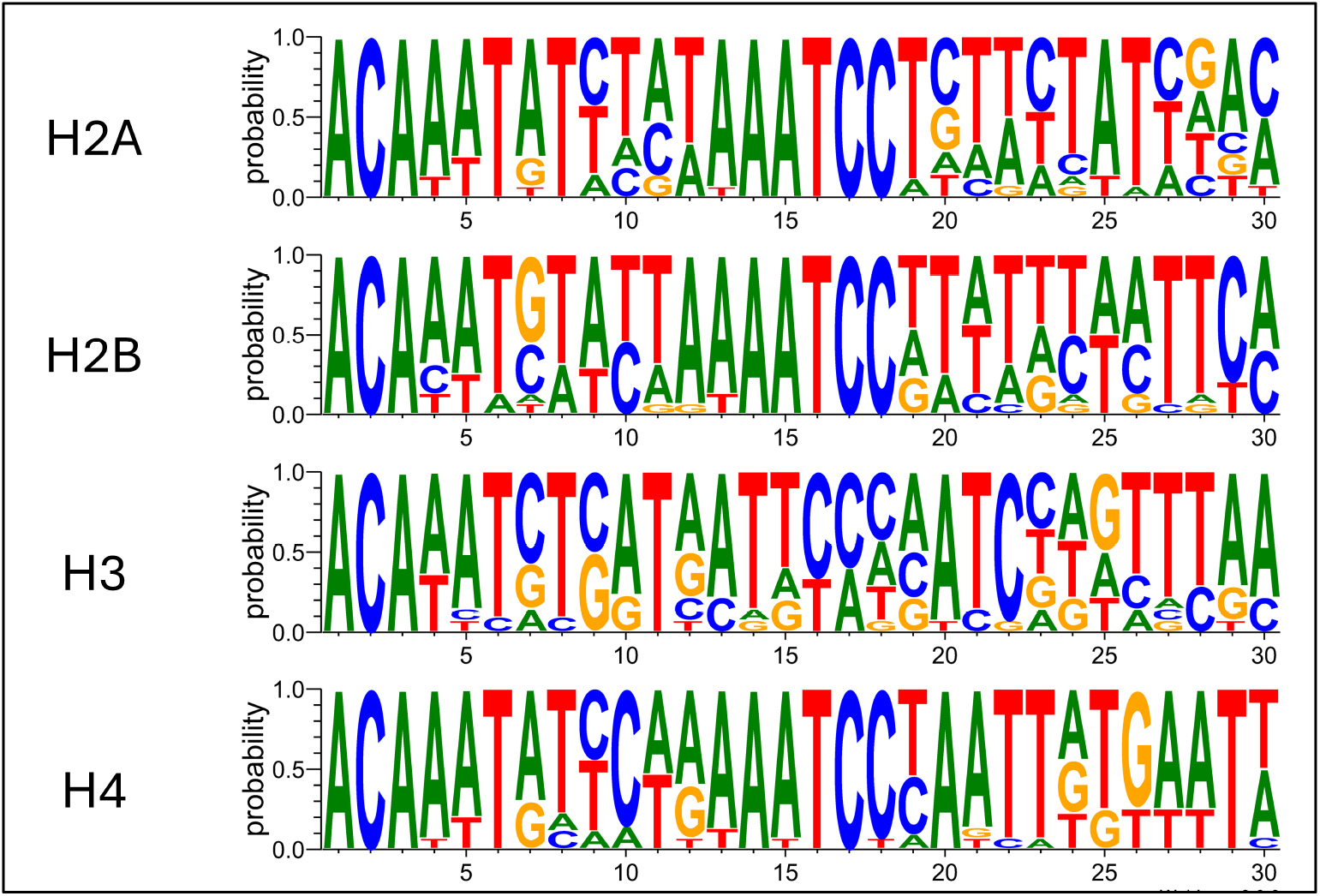
*C. elegans* histone 3’ end sequences. Sequence logo of nucleotides downstream of stem loop for replication-dependent *C. elegans* histones encoding H2A, H2B, H3, and H4. Sequence information were obtained from Wormbase with sequence logo generated by WebLogo3 (37, 38).

To test the strain, we knocked down *cdl-1* (cell death lethal) via RNAi which encodes the homologue of the human SLBP (stem loop binding protein) that serves as the principal protein that facilitates histone 3’ mRNA processing (2). In worms fed with empty vector (EV) RNAi, GFP was mostly absent with faint signals observed in the posterior intestinal cells (**Fig. 1c**). This reflects normal cleavage of *his-6* that results in the loss of the distal GFP transcript; the faint expression of GFP signal in wildtype worms might reflect the overexpressed nature of the reporter. In worms with RNAi knockdown of *cdl-1*, we observed strong activation of GFP throughout the intestinal nuclei, reflecting cleavage failure resulting in transcriptional read-through and activation of the misprocessing reporter (**Fig. 1c**). The nuclear expression of GFP suggests that the construct is expressed as a fusion to the HIS-6 protein, as the GFP variant used does not express a nuclear localization signal. To validate that the increase in GFP fluorescence reflects misprocessing of the endogenous histone transcripts, we designed a pair of primers to distinguish total and misprocessed histone transcripts in response to *cdl-1* knockdown (**Fig. 1d**). RNAi knockdown of *cdl-1* had no effect on total levels of *his-6*, but significantly increased the abundance of misprocessed transcripts (**Fig. 1e**). We also express the two forms of histone transcripts as a ratio of misprocessed / total to normalize for potential fluctuations in the overall histone transcript abundance across samples and found that *cdl-1* depletion consistently increased the misprocessed / total ratio. Given that H3 variants had the greatest divergence in putative HDE motif, we then extended the measurement to genes encoding other histone variants and found the *cdl-1* depletion increased the ratio of misprocessed transcripts of *his-16* (H2A), *his-11* (H2B), and *his-26* (H4) (**Fig. 1e**). Overall, we constructed and validated an *in vivo* histone misprocessing reporter in *C. elegans* that provides a visual biomarker for post-transcriptional histone 3’ cleavage failure.

### Identification of novel histone 3’ processing regulators

To identify genetic regulators of histone 3’ processing, we next performed a genome-wide RNAi screen to identify genes that when knocked down resulted in activation of the histone misprocessing reporter (**Fig. 2a**). We screened ∼19,000 genes from the MRC genomic and ORFeome RNAi library and identified 30 genes that when silenced by RNAi consistently activate GFP expression of the misprocessing reporter (**Fig. 2b, S3**). All 30 genes were found to be essential as RNAi depletion caused either developmental arrest or sterility, consistent with indispensable requirement of correctly cleaved histones for viability. Interestingly, we found that many of the identified genes function within the rRNA biogenesis pathway or encode ribosome subunits (**Fig. 2c**). We also found that knockdown of *ints-4* encoding part of the catalytic module of the Integrator complex results in histone 3’ misprocessing (**Fig. 2c**). This is consistent a previous study showing that inhibition of human Integrator blocks 3’ processing of replication-dependent histones (39, 40).

**Figure 2.**
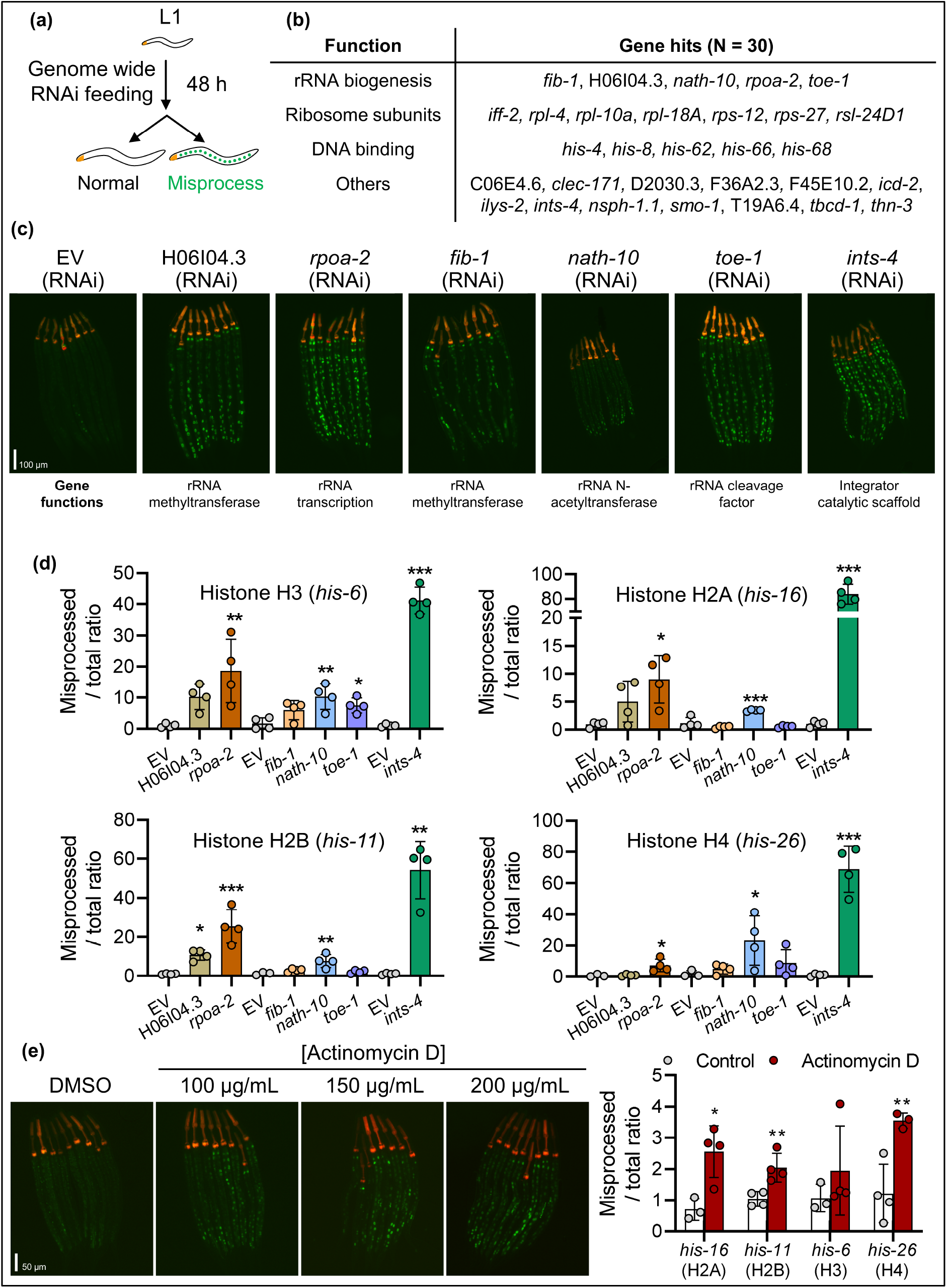
Disruption of rRNA biosynthesis causes histone mRNA 3’ misprocessing. **a)** Outline of the genome-wide RNAi screen to identify gene knockdowns that activate the histone misprocessing reporter. **b)** Categories of verified gene knockdowns that activate the histone misprocessing reporter identified by the genome-wide RNAi screen. **c)** Representative fluorescent micrographs of the histone misprocessing reporter fed with genes encoding the rRNA biosynthesis pathway (H06I04.3, *rpoa-2*, *fib-1*, *nath-10*, *toe-1*) and the Integrator complex (*ints-4*). Eight worms are shown in each image; the scale bar is 100 µm. Functions of each gene are indicated below each image. **d)** Relative levels of misprocessed / total ratio of histone variant genes after RNAi knockdown of the corresponding genes. Multiple EV controls are graphed, as the samples were obtained from multiple experiments. Bar graphs show mean ± STDEV with N = 3-4 biological replicates measured for each condition. One-way ANOVA was used for comparison of H06I04.3, *rpoa-2*, *fib-1*, *nath-10*, and *toe-1*, and student’s t-test was used for comparison of *ints-4.* **e)** Representative fluorescent micrograph of the histone misprocessing reporter treated with DMSO (1% control) or increasing concentrations of Actinomycin D. Eight worms are shown in each image; the scale bar is 50 µm. Bar graph shows relative levels of misprocessed / total ratio of histone variant genes after treatment with 1% DMSO or 200 µg/mL of Actinomycin D as determined by qPCR. Student’s t-test was used for statistical comparison. Statistical significance are denoted by *P<0.05, **P<0.01, ***P<0.001 compared to EV in **d** and control in **e**.

**Figure S3.**
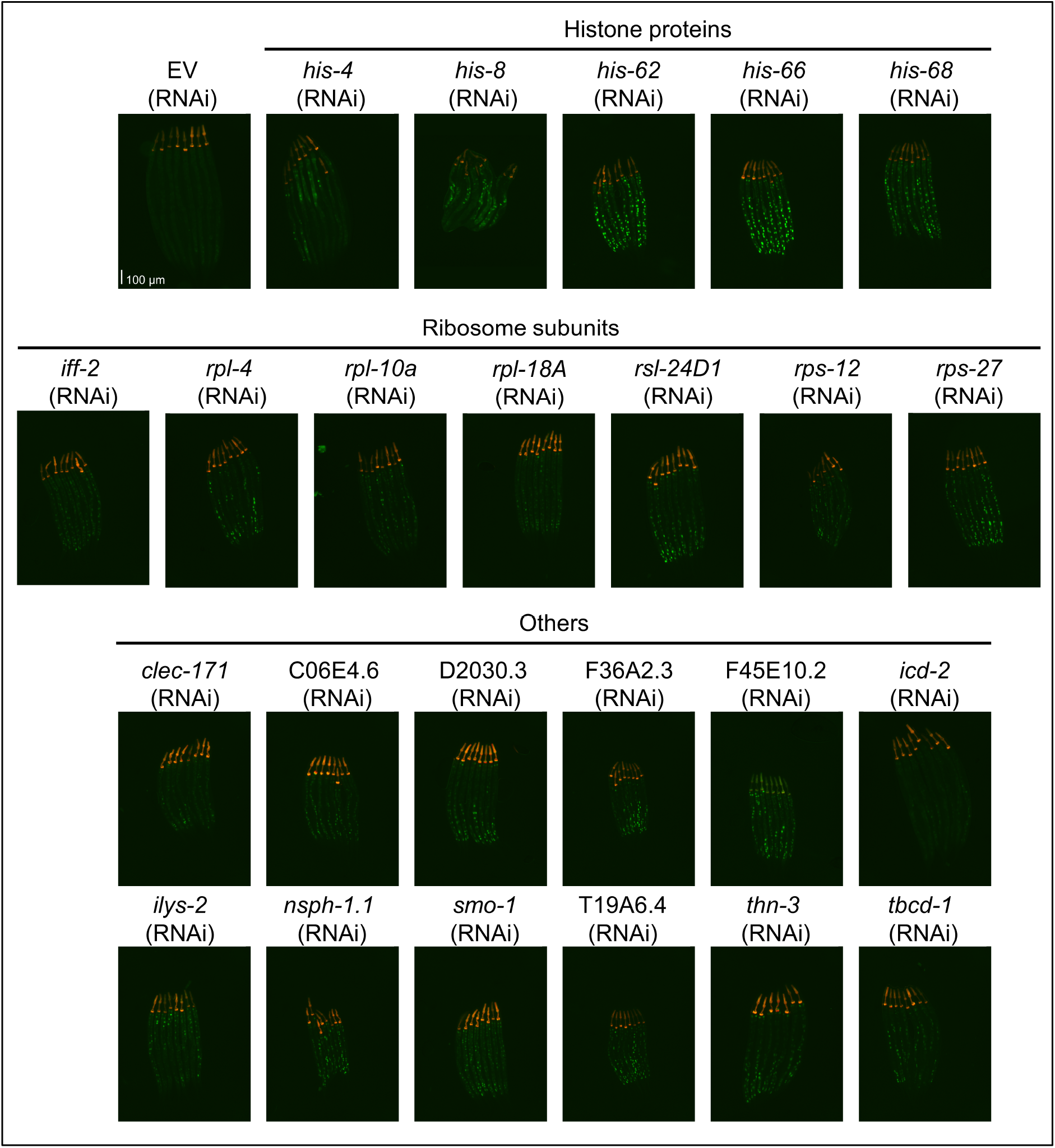
RNAi gene knockdowns that induced histone misprocessing. Representative fluorescent micrographs showing all confirmed hits from the genome-wide RNAi screen that activate the histone misprocessing reporter. Scale bar of 100 µm is shown in EV (RNAi).

We decided to focus on genes functioning in rRNA biogenesis as defects in this pathway have not previously been linked to histone misprocessing. We used qPCR to verify that RNAi knockdown of *rpoa-2*, *nath-10*, and *toe-1* resulted in a significant increase in the ratio of misprocessed *his-6* transcripts (**Fig. 2d**). RNAi knockdown of H06I04.3 (∼10-fold increase, P=0.07) and *fib-1* (∼3.5-fold, P=0.06) also elevated the misprocessed ratio but was not considered statistically significant. We next measured the misprocessed ratio of the other three histone variants and found that across the five rRNA biogenesis genes examined, only knockdown of *rpoa-2* encoding the largest subunit of RNA Pol I resulted in consistent misprocessing of all four histone variants (**Fig. 2d**). For all conditions assessed, total levels of the four histone variant transcripts were not elevated, with select gene knockdowns resulting in a significant decrease in total histone levels (**Fig. S4a**). We decided to focus on *rpoa-2* for the remainder of the study.

We next tested whether histone misprocessing can be triggered by inhibition of rRNA transcription by Actinomycin D. We found that similar to *rpoa-2* knockdown, treatment with Actinomycin D resulted in the activation of the histone misprocessing reporter and increased misprocessed histone transcripts (**Fig. 2e, S4b**). However, the degree of histone misprocessing from Actinomycin D was less pronounced compared to *rpoa-2* RNAi, which may be attributed to the general inefficiency of chemical uptake in worms (41). Given that defects in *rpoa-2* has been shown to decrease the transcription of all three major rRNA transcripts (25), we next designed RNAi clones to individually knockdown 5.8s (*rrn-2.1*), 18s (*rrn-1.1*), and 28s (*rrn-3.1*) rRNA transcripts. Knockdown of individual rRNA did not result in activation of the misprocessing reporter (**Fig. S4c**), suggesting that the effects of *rpoa-2* knockdown on histone misprocessing are unlikely to be caused by depletion of individual rRNA transcripts. Together, the results here identified several new regulators of histone post-transcriptional 3’ cleavage and implicate a central requirement of rRNA biogenesis in this process.

**Figure S4.**
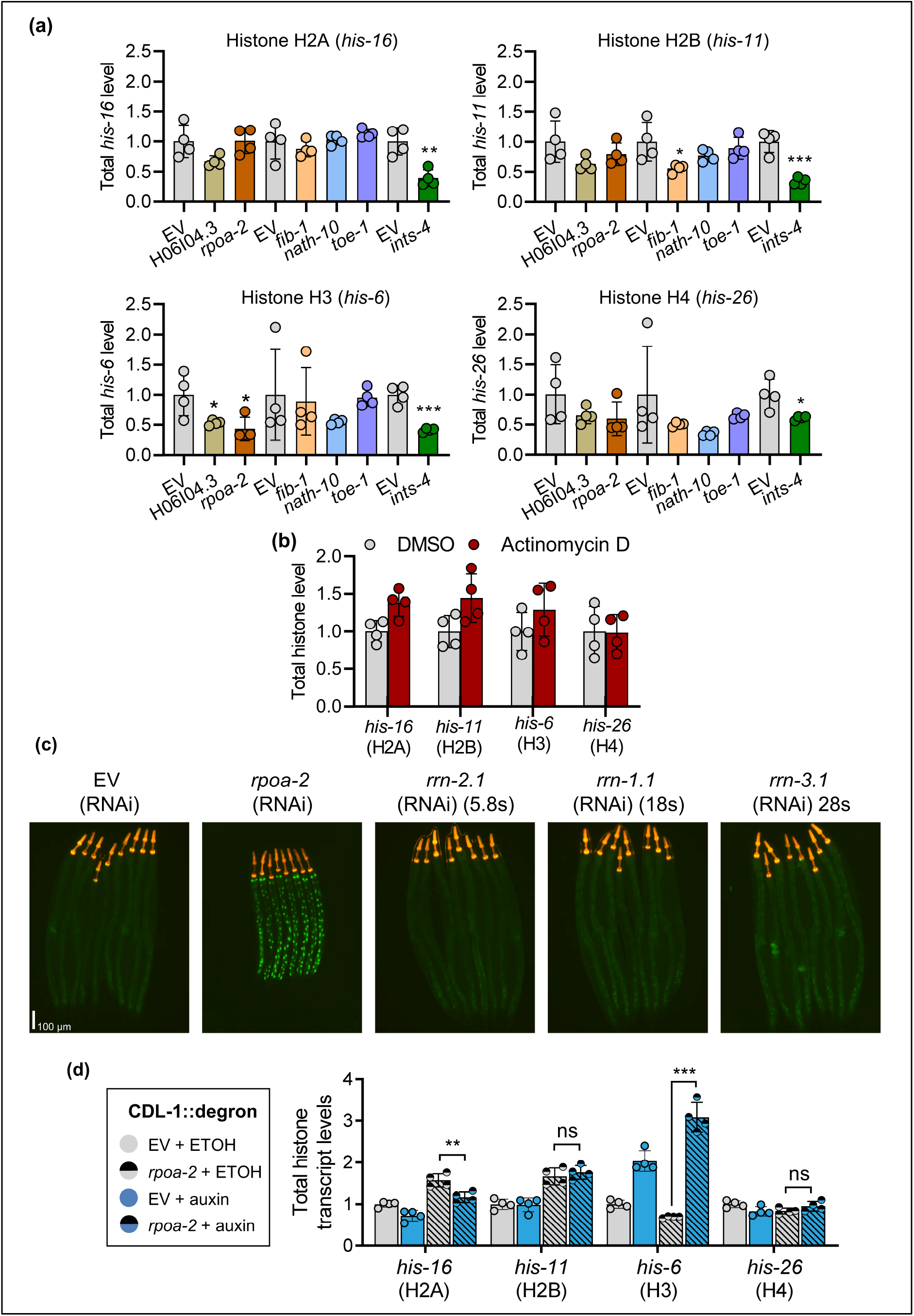
Disruption of rRNA biosynthesis minimally impacts total histone levels. **a)** Relative levels of total transcripts encoding histone H3 (*his-6*), histone H2A (*his-16*), histone H2B (*his-11*), and histone H3 (*his-26*) after RNAi knockdown of H06I04.3, *rpoa-2*, *fib-1*, *nath-10*, *toe-1*, and *ints-4.* **b)** Total levels of histone variant mRNA after treatment with 1% DMSO or 200 µg/mL of Actinomycin D. **c)** Effects of knocking down individual rRNA transcripts on histone misprocessing reporter. *rpoa-2* RNAi is used as a positive control. **d)** Total levels of histone variant mRNA genes in CDL-1::degron strain expressing *eft-3p*::TIR1 and *mex-5p*::TIR1 fed with EV or *rpoa-2* RNAi at the L1 stage on ETOH or 1 mM auxin plates for 48 hours. Bar graphs representing qPCR data show mean ± STDEV with N = 4 biological replicates measured for each condition *P<0.05 **P<0.01 and ***P<0.001 as determined by one-way ANOVA in **a** and two-way ANOVA in **d**. Student’s t-test was used for **b** and for comparison of EV to *ints-4* in **a**.

### rpoa-2 potentiates CDL-1 histone 3’ processing

To test whether *rpoa-2* knockdown influence CDL-1 dependent histone 3’ processing, we first used CRISPR to insert an AID*::3xflag::mNeonGreen epitope to the C-terminal end of the *cdl-1* locus to permit visualization and auxin induced degradation of the endogenous CDL-1 protein (**Fig. 3a**). Fluorescent mNG expression of CDL-1 was visible in the hypodermal and intestinal nuclei in L2 worms, with strong intestinal and germline expression observed at L4 (**Fig. 3b**). At adulthood, somatic expression of CDL-1 is markedly reduced with germline showing the highest expression (**Fig. 3b**). As a *cdl-1* loss of function mutants are not viable, we next introduced the TIR1 F-box protein into this strain expressed under the *eft-3* and *mex-5* promoter to enable simultaneous somatic and germline depletion of CDL-1 with auxin (this strain is referred to as CDL-1::degron) (42). This is accomplished by using PCR to genotype and isolate worms expressing homozygous insertion of the degron tag and TIR1 protein after mating (**Fig. 3c**) (42). To evaluate for potential genetic interactions between *rpoa-2* and CDL-1, we knocked down *rpoa-2* via RNAi and depleted CDL-1 via auxin to determine the effects of single or double knockdown on histone misprocessing. Surprisingly, we found that apart from *his-6*, *rpoa-2* knockdown led to a further increase in misprocessed histone transcripts in worms with CDL-1 co-depleted via auxin (**Fig. 3d**). We observed a decrease in misprocessed *his-6* ratio in the double depletion worms that was attributed to a 4.5-fold increase in total *his-6* levels in this group, an effect that was not observed in the other histone variants (**Fig. S4d**). Together, the co-depletion experiments suggest that *rpoa-2* and CDL-1 may act in an independent or a partially redundant pathway to influence histone 3’ processing.

**Figure 3.**
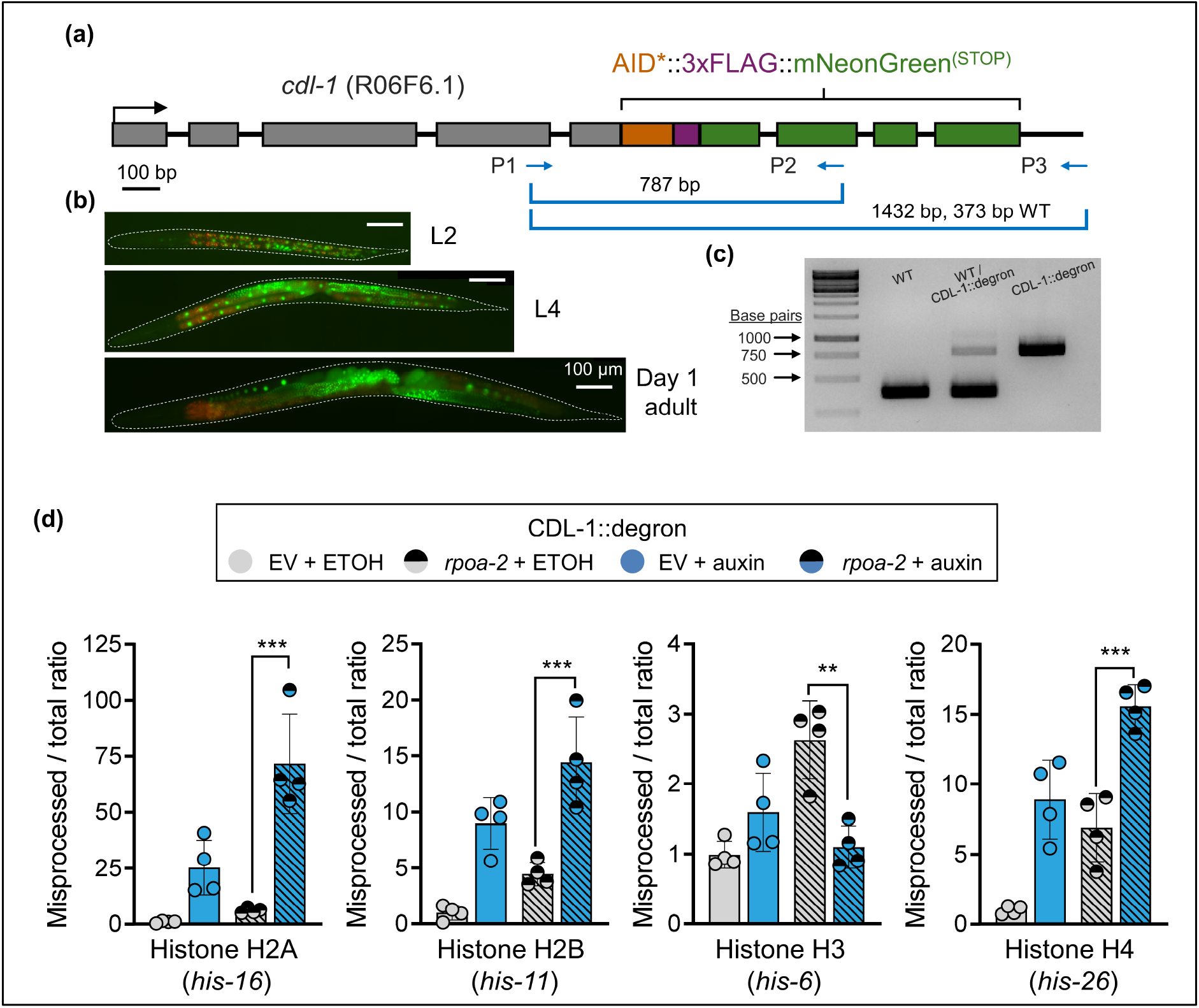
Co-depletion of *rpoa-2* and CDL-1 exacerbates histone mRNA misprocessing. **a)** Schematic of CDL-1::degron strain generated by CRISPR insertion of AID*::3xflag::mNG in the C-terminal of the *cdl-1* locus. Primers used to genotype the insertion are illustrated and listed in Table S2. **b)** Expression of CDL-1 fused with mNG at different developmental stages. Composite images shown were created by merging images taken with the GFP and DAPI channels to distinguish autofluorescence (orange) from CDL-1::mNG signals. **c)** PCR gel showing corresponding DNA fragments amplified from wildtype N2, heterozygote, and homozygote AID*::3xflag::mNG inserted animals using the P1-3 triple primers. Fluorescent micrographs show the expression pattern of CDL-1 at larval and adult stages as visualized by the mNG signal. Scalebar is 100 µm. **d)** Relative levels of misprocessed / total ratio of histone variant genes in CDL-1::degron strain expressing *eft-3p*::TIR1 and *mex-5p*::TIR1 fed with EV or *rpoa-2* RNAi at the L1 stage on ETOH or 1 mM auxin plates for 48 hours. Bar graphs show mean ± STDEV with N = 4 biological replicates measured for each condition **P<0.01 and ***P<0.001 as determined by two-way ANOVA.

To further explore this interaction, we perform mRNA-sequencing to survey the effects of *rpoa-2* and CDL-1 depletion have on the transcriptome (**Fig. 4a**). Principal component analysis (PCA) using all normalized gene counts demonstrated that *rpoa-2* knockdown and CDL-1 depletion clustered closely together, but the combined *rpoa-2* and CDL-1 co-depletion led to a separation from both single depletion effects (**Fig. 4b**). This suggests that combining *rpoa-2* and CDL-1 depletion produces a distinct transcriptome, which was also supported by the Euclidean distance analysis (**Fig. 4b**). We first compared the transcriptome profile between single *rpoa-2* knockdown and CDL-1 depletion. Through Pearson correlation analysis, we found that for genes that were differentially regulated by >2-fold by CDL-1 depletion, there was a striking similarity in the expression profile in *rpoa-2* knockdown worms with a correlation value of 0.84 (**Fig. 4c**). Of the 2,245 CDL-1 depletion-induced differentially expressed genes, ∼94% of the genes were altered in the same direction by *rpoa-2* depletion (**Fig. 4c**). We also inverted the analysis for the 4,339 genes differentially expressed by >2-fold by *rpoa-2* knockdown and observed a consistent correlation value of 0.83 compared to the expression changes induced by CDL-1 depletion, with ∼95% of the genes altered in the same direction (**Fig. S5a**). This indicated that both *rpoa-2* and CDL-1 regulate a similar set of target genes. To better understand this relationship, we first focused on examining transcripts that were down-regulated and found that the majority of the genes (826/974) that were decreased by >2-fold after CDL-1 depletion were co-regulated by *rpoa-2* (**Fig. 4d**). Enrichment analysis of the shared down-regulated targets shows that they cluster strongly to functions within the cell cycle and division regulation; this suggests that *rpoa-2* was also required for regulating CDL-1 dependent cell cycle activities. Examining genes up-regulated by *rpoa-2* knockdown and CDL-1 depletion also yielded significant overlap in shared regulatory targets (713/1,271). However, in contrast to the highly specific enrichment to cell cycle regulators as observed for down-regulated genes, enrichment analysis of the 713 shared genes shows clustering to a limited number of processes related to cuticle molting and immune response pathways (**Fig. S5b**).

**Figure 4.**
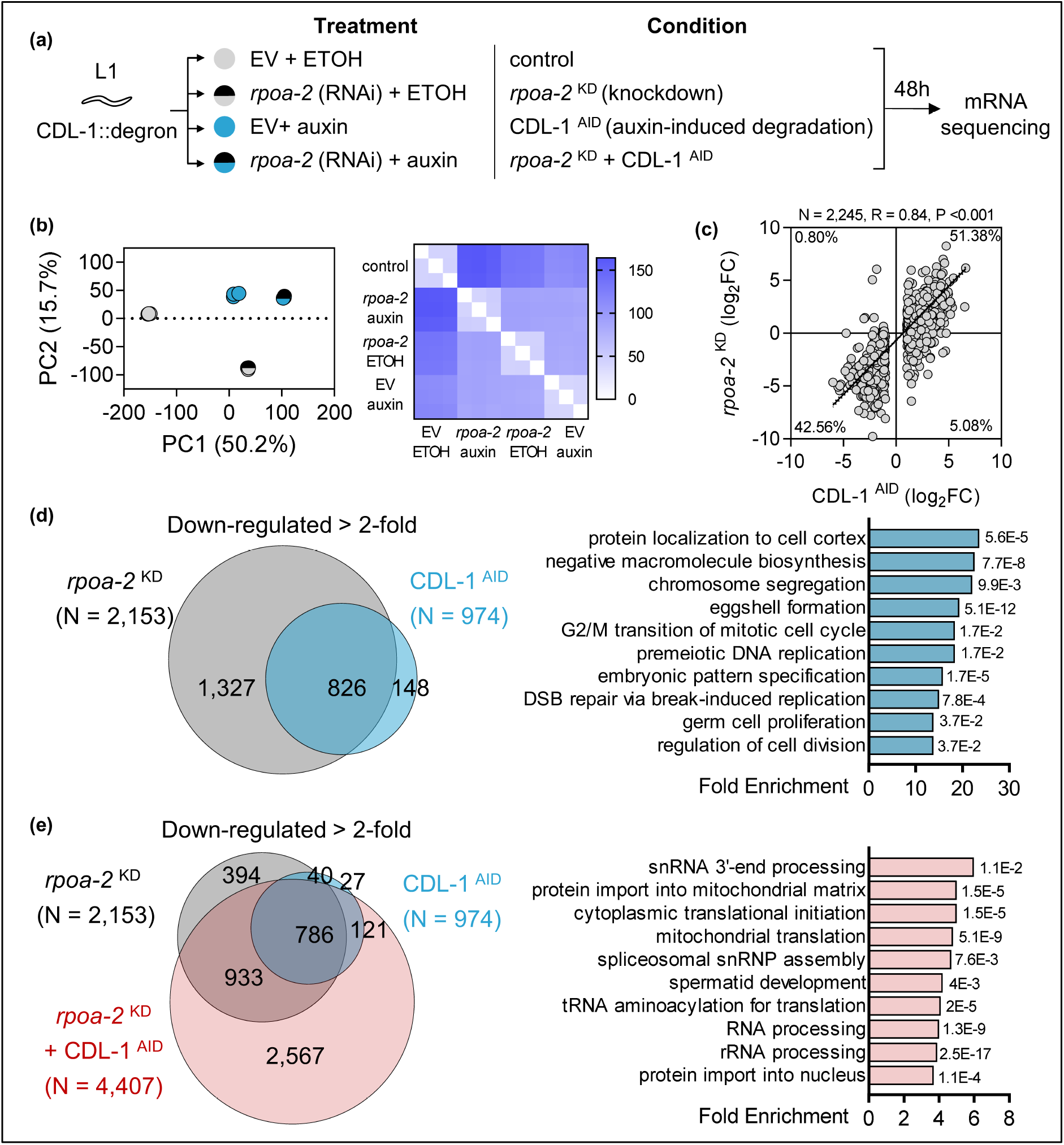
CDL-1 and *rpoa-2* overlap in transcriptome regulation. **a)** Workflow of treatment conditions for mRNA sequencing. **b)** Principal component analysis and heatmap of Euclidean sample distance of the four mRNA sequencing experimental conditions. **c)** Correlation analysis of log_2_ fold change between CDL-1 degron depletion (CDL-1 ^AID^) and *rpoa-2* RNAi knockdown (*rpoa-2* ^KD^) for 2,245 differentially expressed genes by more than 2-fold detected in CDL-1 depleted worms. Pearson’s correlation was used to test statistical significance. **d)** Venn diagram illustrating the number of genes down-regulated by >2-fold overlapping between CDL-1 ^AID^ and *rpoa-2* ^KD^. Cluster analysis of biological processes enriched by the 826 shared genes. **e)** Venn diagram illustrating the number of genes down-regulated by >2-fold overlapping between CDL-1 ^AID^, *rpoa-2* ^KD^, and CDL-1 ^AID^ + *rpoa-2* ^KD^. Cluster analysis of biological processes enriched by the 2,567 genes uniquely down-regulated in the CDL-1 ^AID^ + *rpoa-2* ^KD^ condition.

A notable observation from the PCA and Euclidean distance analysis is that double depletion of *rpoa-2* and CDL-1 led to a unique transcriptome profile (**Fig. 4b**). We first examined the shared down-regulated genes between single or double *rpoa-2* and CDL-1 depletion and found that there were 2,567 genes that were down-regulated only when both *rpoa-2* and CDL-1 were co-depleted (**Fig. 4e**). Enrichment analysis of these 2,567 down-regulated transcripts shows that they encode genes that are broadly involved in RNA processing and localized to the nucleus or the mitochondria. Notably, genes encoding subunits of the Integrator complex were down-regulated and clustered under the GO term snRNA 3’-end processing, where double depletion caused >2-fold down-regulation in 7 Integrator subunit genes, which was not seen when *rpoa-2* or CDL-1 were individually depleted (**Table S2**). We also found that double *rpoa-2* and CDL-1 depletion caused >2-fold down-regulation in 9 out of 30 genes from the genome-wide RNAi list. This suggests that co-disruption of *rpoa-2* and CDL-1 caused transcriptional decrease in many identified histone 3’ processing regulators, which was not observed when *rpoa-2* or CDL-1 was depleted individually. This in part could explain why double *rpoa-2* and CDL-1 depletion caused a synergistic increase in the abundance of misprocessed histone transcripts (**Fig. 3d**). Double *rpoa-2* and CDL-1 depletion also resulted in the exclusive up-regulation of 638 genes, however, these genes enrich to a broad range of processes, including oxygen response, secondary messenger synthesis, and cilium organization (**Fig. S5c**).

Together, transcriptome profiling indicates that *rpoa-2* and CDL-1 are both required to positively regulate the expression of cell cycle regulating genes. Additionally, *rpoa-2* and CDL-1 cooperatively control a subset of genes that are broadly involved in RNA metabolism, including genes that were found to regulate histone 3’ processing such as those functioning in rRNA processing and the Integrator complex.

**Figure S5.**
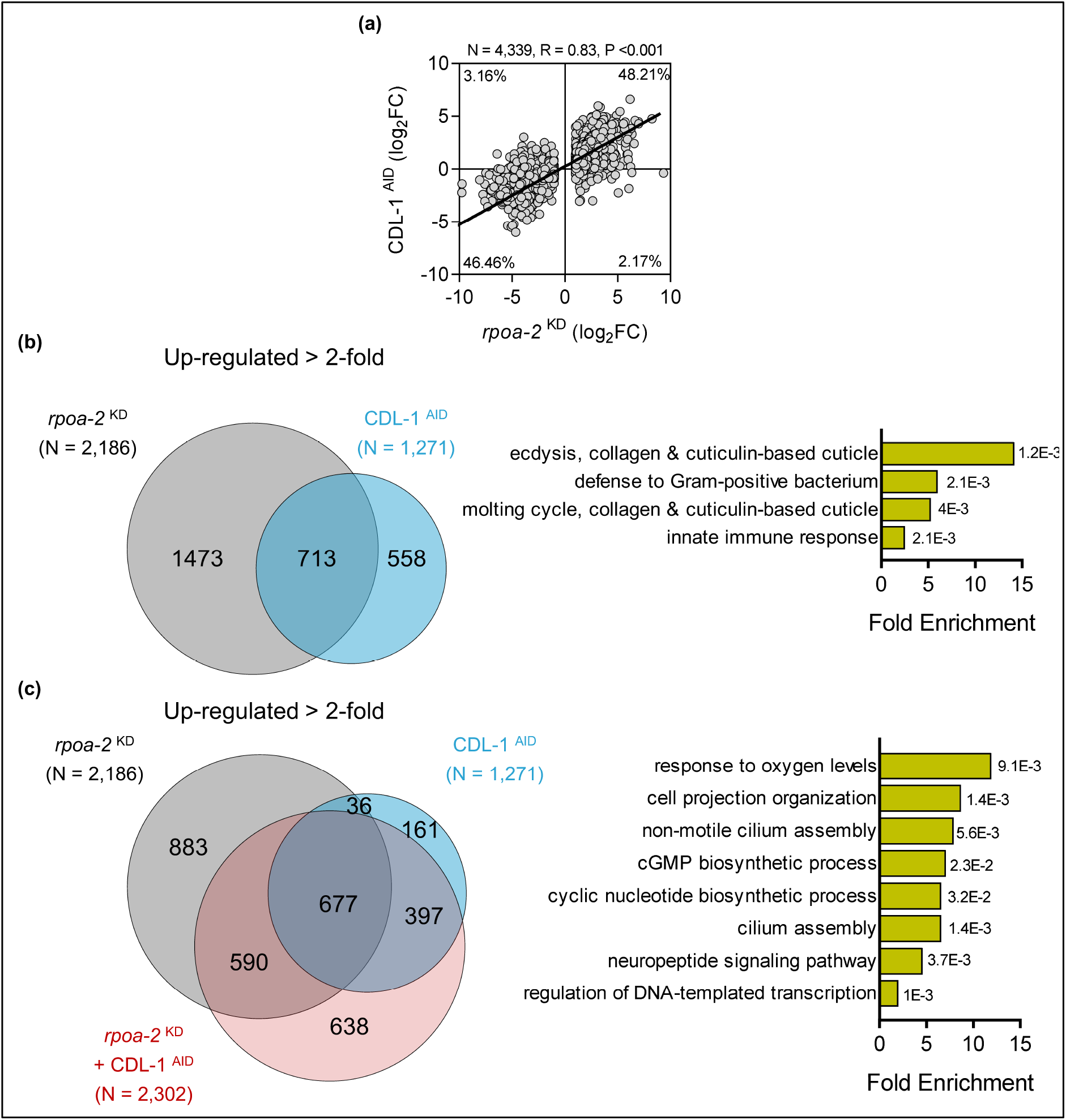
CDL-1 and *rpoa-2* overlap in gene up-regulation. **a)** Correlation analysis of log_2_ fold change between *rpoa-2* RNAi knockdown (*rpoa-2* ^KD^) and CDL-1 degron depletion (CDL-1 ^AID^) for 4,339 differentially expressed genes by more than 2-fold detected in *rpoa-2* knocked down worms. Pearson’s correlation was used to test statistical significance. **b)** Venn diagram illustrating the number of genes up-regulated by >2-fold overlapping between CDL-1 ^AID^ and *rpoa-2* ^KD^. Cluster analysis of biological processes enriched by the 713 shared genes. **c)** Venn diagram illustrating the number of genes up-regulated by >2-fold overlapping between CDL-1 ^AID^, *rpoa-2* ^KD^, and CDL-1 ^AID^ + *rpoa-2* ^KD^. Cluster analysis of biological processes enriched by the 638 genes uniquely up-regulated in the CDL-1 ^AID^ + *rpoa-2* ^KD^ condition.

### rpoa-2 depletion induces aberrant cell cycle progression

Our transcriptomic data suggests that *rpoa-2* is a key positive regulator of genes functioning in various cell cycle processes ranging from DNA replication, cell cycle transitions, to DNA damage repair (**Fig. 4**). To evaluate how *rpoa-2* influences the cell cycle, we employed a strain of worm expressing a single copy of the S-phase marker Proliferating Cell Nuclear Antigen homologue protein PCN-1 fused to GFP, along with a codon-optimized human DNA Helicase B (DHB) fragment fused to mKate2 that serves as a CDK activity sensor (43). This strain allows for *in vivo* analysis of cell cycle progression based on PCN-1 and DHB nuclear localization (43). When cells are in the G_0_/G_1_ phase, low CDK activity results in the nuclear localization of the DHB::mKate2 reporter; as cells transition to S and G_2_/M phase, there is a progressive decline in nuclear DHB::mKate2 signal as CDK phosphorylation activates the nuclear export signal to promote cytoplasmic expression (43, 44). We focused on the expression of these markers in the intestinal nuclei as this is the tissue we observed the highest expression of the histone misprocessing reporter (**Fig. 2c**).

RNAi depletion of *rpoa-2* did not cause changes in total PCN-1::GFP fluorescence; however, the relative proportion of nuclear PCN-1::GFP in the intestinal cells was significantly increased by 76% (EV: 15.57% ± 6.31 vs. *rpoa-2*: 27.54% ± 5.65; **Fig. 5a**). We then examined the pattern of DHB fluorescence where we found that under control conditions, DHB was strictly expressed in the intestinal nuclei, indicating low CDK activity that is consistent with the end of the endoreplication cycle in L4 worms. However, in *rpoa-2* knocked down worms, DHB signal was visibly diffused in the intestinal cells with a significant increase in the cytoplasmic to nuclear ratio, indicating the retention of CDK activity in promoting DHB nuclear export (**Fig. 5b**).

**Figure 5.**
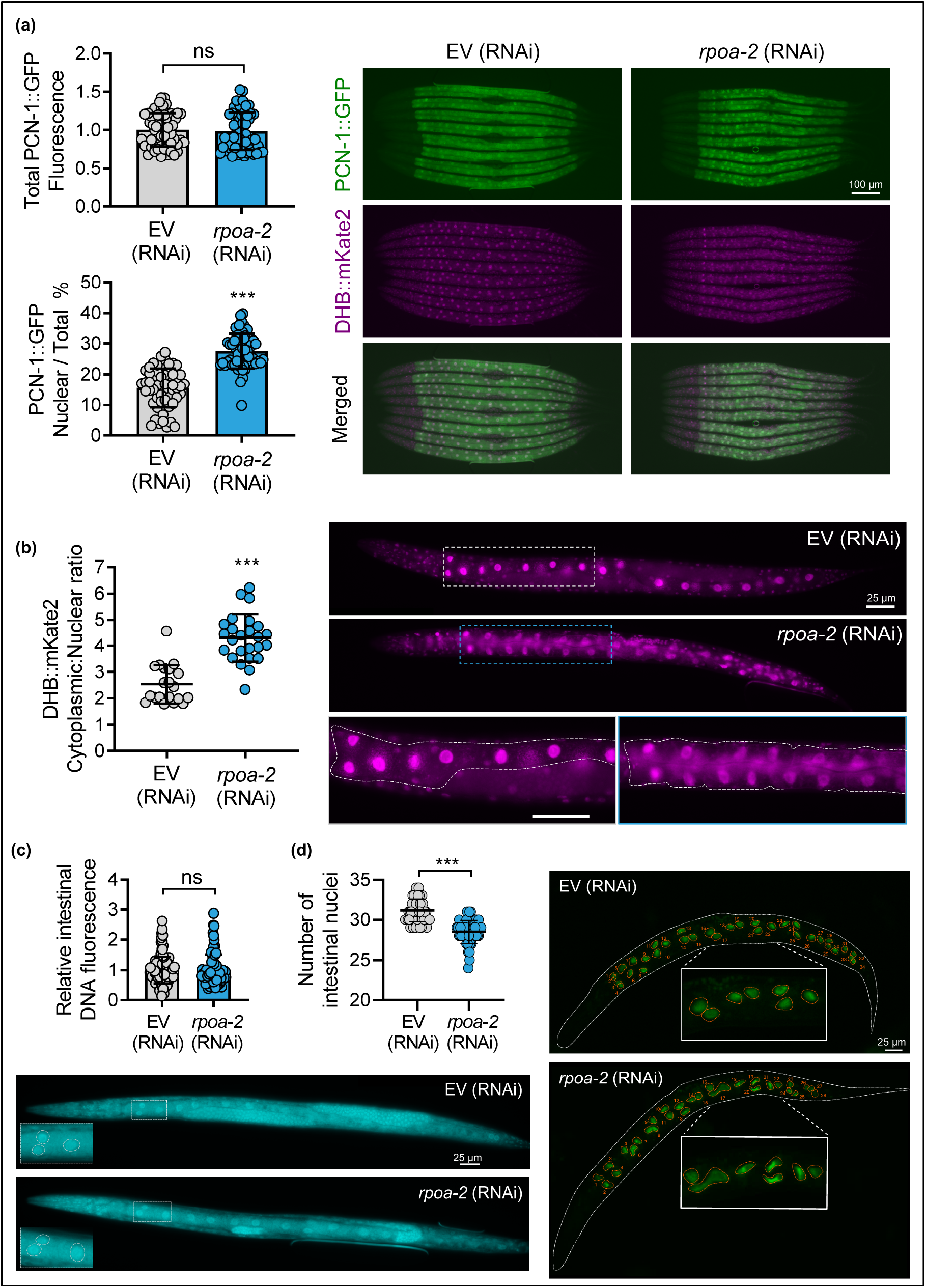
Depletion of *rpoa-2* results in deregulated cell cycle markers. **a)** Quantification of total PCN-1 GFP fluorescence and nuclear/total PCN-1 GFP ratio in worms fed with EV or *rpoa-2* RNAi. Representative fluorescent micrographs showing PCN-1::GFP, DHB::mKate2, and merged composites of the dual reporter with eight worms per image. N = 56 worms quantified from 2 independent trials for each condition. **b)** Quantification of cytoplasmic/nuclear ratio of DHB::mKate2 in worms fed with EV or *rpoa-2* RNAi. Representative image showing whole worm with box sections magnified to illustrate DHB::mKate2 signal patterns. The intestine is outlined in white. N = 20 (EV) or 25 (*rpoa-2*) worms quantified individually from two trials. **c)** Quantification of total intestinal DNA fluorescence in N2 worms fed with EV or *rpoa-2* RNAi. N = 124 nuclei from 20 worms and N = 97 nuclei from 18 worms quantified from EV and *rpoa-2* RNAi, respectively, from 2 independent trials. Representative image showing whole worm staining with box sections magnified to illustrate examples of intestinal nuclei quantified. **d)** Quantification of the number of intestinal nuclei as visualized by GFP tagged histone H2B (*his-48*) in worms fed with EV or *rpoa-2* RNAi. Intestinal nuclei are outlined and numbered in orange. Box sections magnified to illustrate the nuclei shape; a total of 51-53 worms were scored per condition combined from two independent trials. For all experiments, worms were fed with corresponding RNAi starting at L1 for 48 hours. ns: not significant, ***P<0.001 as determined by Student’s t-test. Scalebar is 100 µm in **a** and 25 µm in **b**, **c**, and **d**.

The increase in PCN-1 nuclear expression and CDK activity prompted us to examine whether *rpoa-2* influences polyploidy in the intestinal cells, which undergoes endomitosis to produce 30-34 nuclei and endoreplication to increase their DNA content as it develops from L1 to the L4 stage (45). We first looked at total intestinal DNA content where quantification of Hoechst-stained intestinal nuclei did not reveal a difference in relative amounts of DNA between *rpoa-2* and control conditions (**Fig. 5c**). However, we found that *rpoa-2* knocked down worms had an average of 28 intestinal nuclei at the L4 stage, compared an average of 31 nuclei in the control (**Fig. 5d**). In addition, we also observed that many *rpoa-2* intestinal nuclei exhibit abnormal elongated and crescent shapes, which have previously been linked to defect in chromosomal segregation associated with mutations to spindle assembly checkpoint, or after DNA damage triggered by feeding on the gram-negative *Rhizobium* microbe that induce genome instability (46–48). Together, the data here show that depletion of *rpoa-2* causes aberrant activation of cell cycle markers and leads to partial defects in endomitosis and abnormally shaped *C. elegans* intestinal nuclei.

### rpoa-2 disruption triggers histone misprocessing via multiple downstream pathways

To explore how rRNA biogenesis is connected to histone processing, we examined the role of known negative regulators of cell cycle progression for their requirement in mediating *rpoa-2* dependent histone misprocessing. We first examined a strain of worm harbouring the *atm-1(gk186)* allele lacking the kinase domain of the DNA damage response apical kinase Ataxia-Telangiectasia-Mutated (49). We compared the expression of genes that showed >2-fold changes in *rpoa-2* knockdown worms to previously published microarray data on the transcriptome of one day old *atm-1(gk186)* mutants and found a significant negative correlation value of −0.59 (**Fig. 6a**) (50). Inverted analysis of 860 genes differentially regulated >2-fold in the *atm-1(gk186)* mutant also showed a similar negative correlation of −0.63 (**Fig. S6a**). This suggests that loss of *atm-1* has an opposite effect on the transcriptome to *rpoa-2* knockdown, with 82.7% of the *rpoa-*2 dependent genes being expressed in the inverse direction in the *atm-1* mutant (**Figure 6a**). We next depleted *rpoa-2* in wildtype and the *atm-1* mutant worms and found that loss of *atm-1* resulted in significant suppression of histone misprocessing in three out of four variants (**Fig. 6b**). In the *atm-1* mutant, knockdown of *rpoa-2* still induced *his-16* misprocessing to the same degree as the wildtype, but completely rescued the misprocessing of *his-6* and *his-26*. (**Fig. 6b**). This result shows that *rpoa-2* knockdown triggers histone misprocessing through *atm-1* in a histone variant specific manner.

**Figure 6.**
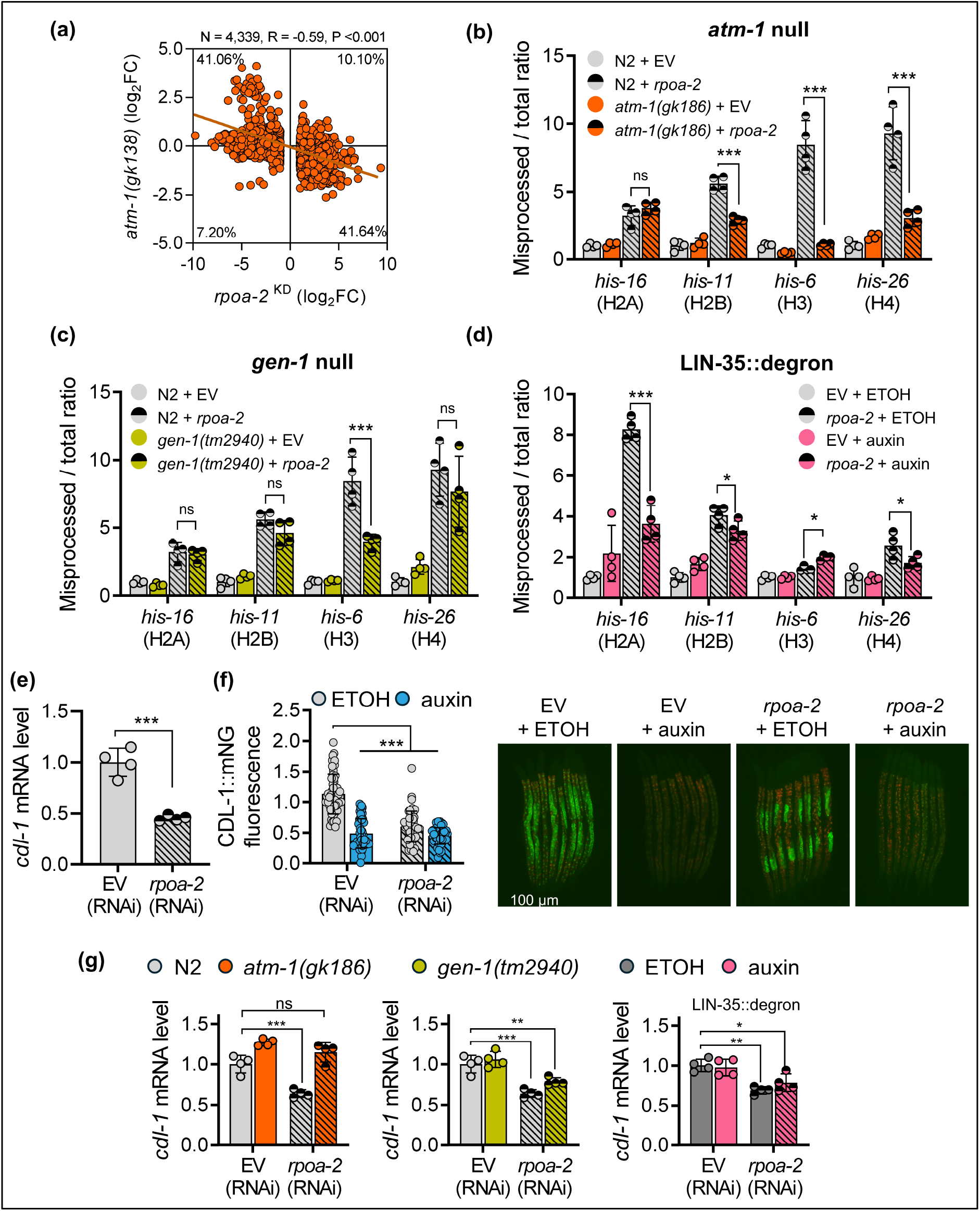
*rpoa-2* signals through cell cycle regulators to induce histone misprocessing. **a)** Correlation analysis of log_2_ fold change between *rpoa-2* RNAi knockdown (*rpoa-2* ^KD^) and *atm-1(gk138)* mutant for 4,339 differentially expressed genes by more than 2-fold detected in *rpoa-2* knockdown worms. Gene expressions for *atm-1(gk138)* were obtained from previously published microarray data and analyzed via limma-voom on the Galaxy server (50, 51). Pearson’s correlation was used to test statistical significance. Relative levels of misprocessed / total ratio of histone variant genes in wildtype N2, **b)** *atm-1(gk138)*, and **c)** *gen-1(tm2940)* null mutants fed with EV or *rpoa-2* RNAi, or **d)** LIN-35::degron strain fed with EV or *rpoa-2* RNAi grown on ethanol or auxin plates. **e)** Relative mRNA levels of *cdl-1* mRNA in N2 worm strains fed with EV or *rpoa-2* RNAi as determined by qPCR. **f)** Quantification of CDL-1::mNG expression in worms fed with EV or *rpoa-2* RNAi grown on ETOH or auxin plates for 48 hours starting from L1. N=48-56 worms measured from 2 independent trials. Representative composite images shown, created by merging images taken with the GFP and DAPI channels to distinguish autofluorescence (orange) from CDL-1::mNG signals. Eight worms are shown in each image. **g)** Relative levels of *cdl-1* mRNA in N2, *cep-1(gk138), gen-1(tm2940)*, or LIN-35 degraded worms relative to control strains fed with EV or *rpoa-2* RNAi. qPCR bar graphs show mean ± STDEV with N = 4 biological replicates measured for each condition; ns: not significant, *P<0.05, **P<0.01, ***P<0.001 as determined by two-way ANOVA in panel **b**, **c**, **d**, **f**, **g** and student’s t-test in **e**.

We next investigated other cell cycle regulators with known roles downstream of *atm-1* including *gen-1* encoding a Holliday junction resolvase, *cep-1* encoding the mammalian p53 transcription factor, and *lin-35* encoding the *C. elegans* retinoblastoma homologue (52–54). Depletion of *rpoa-2* in the *cep-1(gk138)* mutant did not rescue the misprocessing of any histone transcripts (**Fig. S6b**). However, we found that in the *gen-1(tm2940)* mutant, misprocessing of *his-6* after *rpoa-2* knockdown was significantly reduced compared to the wildtype (**Fig. 6c**). To explore *lin-35*, we used a CRISPR generated protein degron depletion strain and found that *rpoa-2* knockdown in ubiquitous LIN-35 depleted worms suppressed misprocessing of *his-16*, *his-11*, and *his-26*, but not *his-6*. (**Fig. 6d, S7a**). These data suggest that *rpoa-2* knockdown signals through multiple downstream regulators independent of *cep-1* to trigger misprocessing of different histone variants.

**Figure S6.**
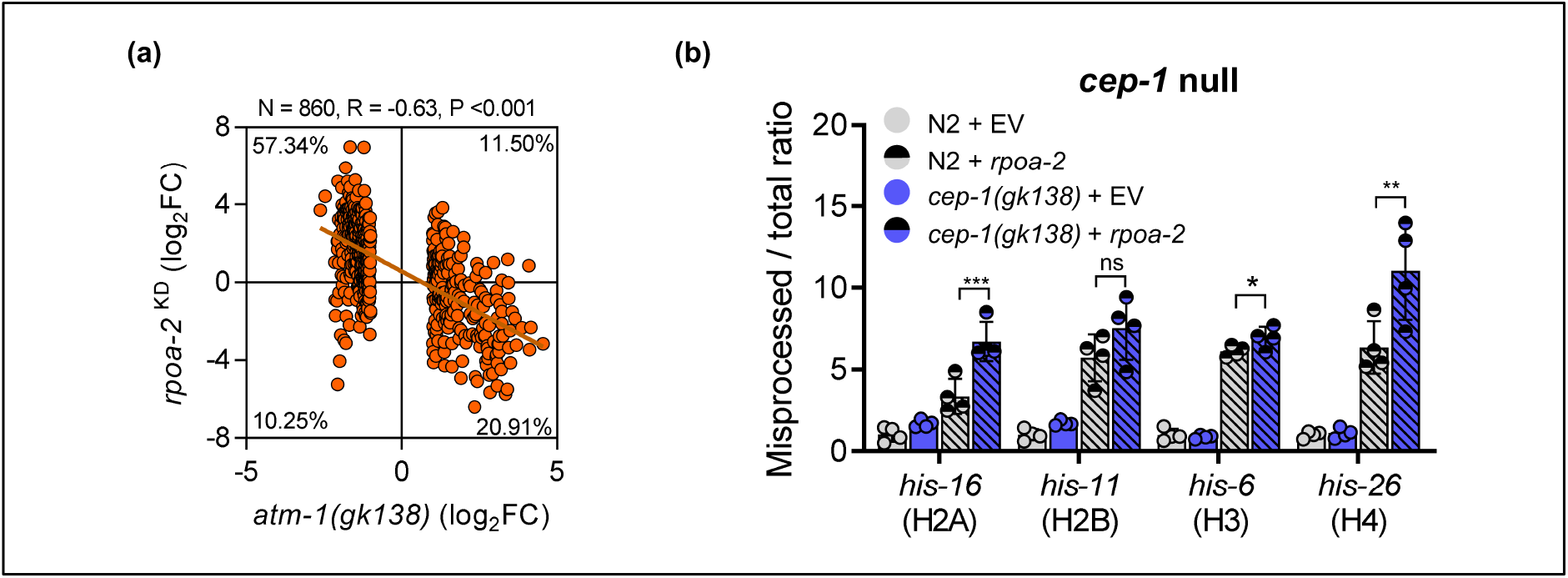
*atm-1* and *cep-1* on *rpoa-2* transcriptome and histone processing. **a)** Correlation analysis of log_2_ fold change between *atm-1(gk138)* and *rpoa-2* knockdown for 860 differentially expressed genes by more than 2-fold detected *atm-1(gk138)* worms (50). **b)** Relative levels of misprocessed / total ratio of histone variant genes in wildtype N2 and *cep-1(gk138)* worms fed with EV or *rpoa-2* RNAi. ns: not significant, *P<0.05, **P<0.01, ***P<0.001 as determined by two-way Anova.

To further define how *rpoa-2* depletion induces histone misprocessing, we re-examined our RNA-seq data and found that *rpoa-2* knockdown caused a 60% down-regulation of the *cdl-1* mRNA, which we confirmed with qPCR and verified independently in the wildtype N2 strain (**Fig. 6e, Fig. S7b**). We then asked if *rpoa-2* knockdown also down-regulate the CDL-1 protein using the degron strain that was fused to mNG. We found the *rpoa-2* depletion significantly reduced overall CDL-1::mNG fluorescence, suggesting that the down-regulation effect occurs at both the transcriptional and translational levels (**Fig. 6f**). Curiously, we observed that *rpoa-2* knockdown caused the strongest reduction in CDL-1::mNG in somatic cells (*i.e.* intestinal nuclei) with mNG signals still retained in the germline. We then used a CRISPR generated endogenously tagged GFP::RPOA-2 strain to evaluate *rpoa-2* RNAi efficiency and found that worms fed with *rpoa-2* RNAi caused a strong reduction in GFP::RPOA-2 signal in somatic cells, but visible germline fluorescence was retained (**Fig. S7c**). As such, we conclude that the retention of CDL-1::mNG fluorescence in the germline after *rpoa-2* knockdown was likely attributed to tissue-specific variability in *rpoa-2* RNAi sensitivity, with reduced efficacy observed in the germline.

**Figure S7.**
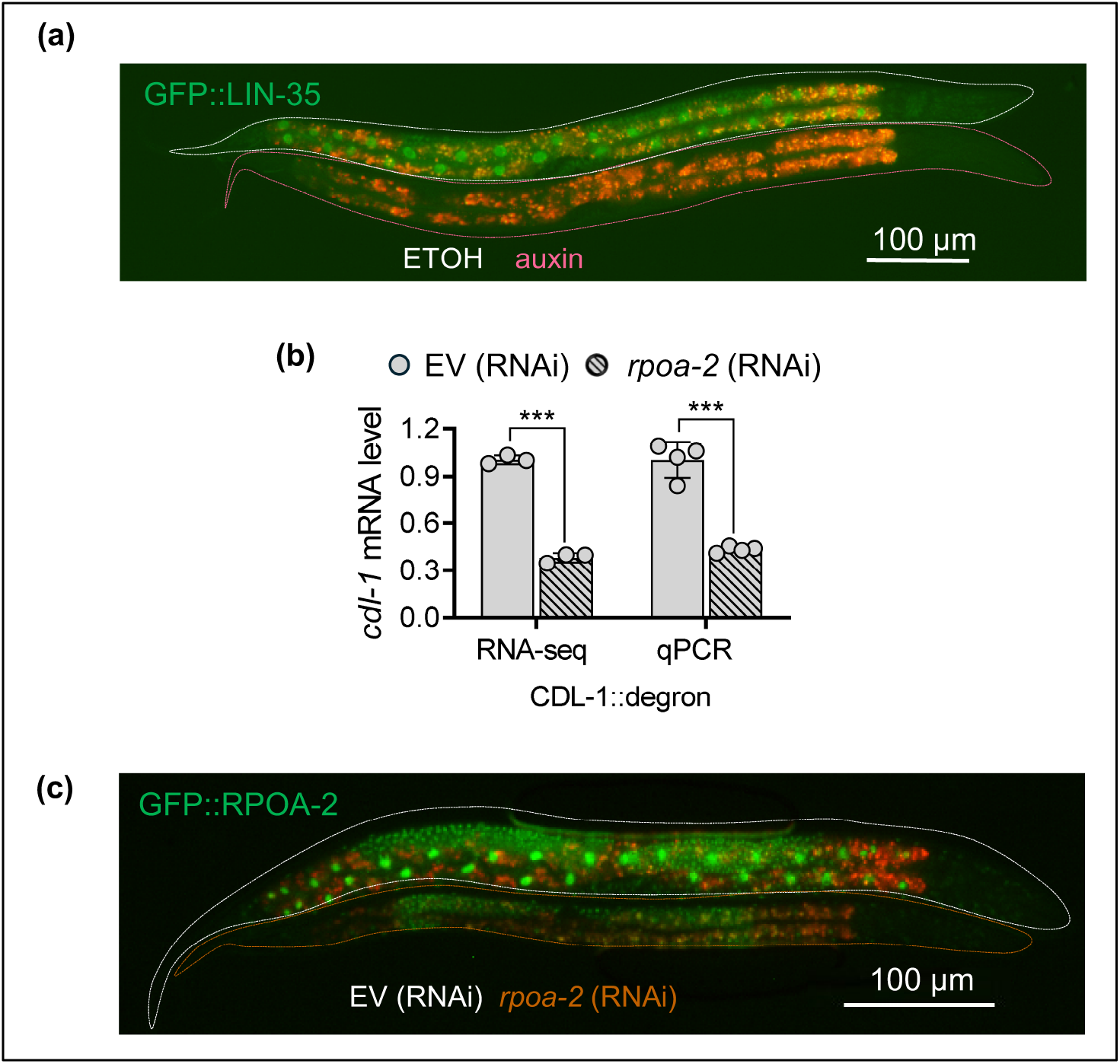
LIN-35 and RPOA-2 target depletion. **a)** Fluorescent micrograph showing effects of auxin treatment on depletion of GFP::LIN-35. **b)** mRNA level of *cdl-1* after *rpoa-2* RNAi as determined by RNA-seq or qPCR in the CDL-1::degron strain. **c)** Fluorescent micrograph showing effects of *rpoa-2* RNAi on endogenous RPOA-2 protein tagged with GFP (22). The composite image shown is created by merging images taken from the GFP and DAPI channel to distinguish intestinal autofluorescence (orange).

To determine if down-regulation of *cdl-1* after *rpoa-2* depletion requires *atm-1*, *gen-1*, or LIN-35, we measured *cdl-1* mRNA in these strains after *rpoa-2* depletion. We found that the *atm-1* mutant completely rescued down-regulation of *cdl-1* after *rpoa-2* knockdown; however, *cdl-1* was still down-regulated by *rpoa-2* RNAi in the *gen-1* mutant or LIN-35 depleted strain to the same degree as observed in the wildtype strain or the control condition respectively (**Fig. 6g**). Overall, these results show that disruption to *rpoa-2* signals through multiple pathways to induce histone misprocessing, by down-regulating *cdl-1* expression in an *atm-1* dependent manner, and via a parallel pathway through *gen-1* and LIN-35 that is independent of *cdl-1* (**Fig. 7**).

**Figure 7.**
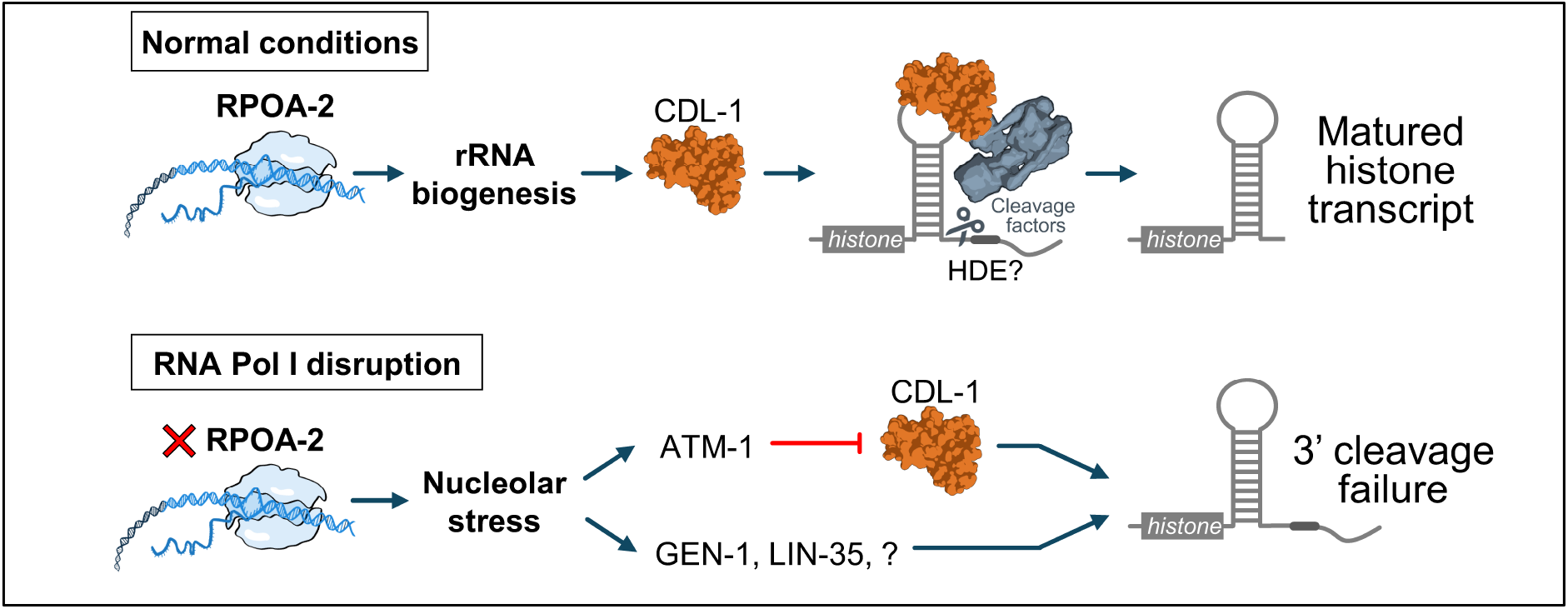
Proposed model of *rpoa-2* dependent histone 3’ processing. Normal RPOA-2 expression regulates rRNA and ribosome biogenesis to support CDL-1 expression for histone 3’ processing. Upon *rpoa-2* depletion, nucleolar stress causes down-regulation of *cdl-1* that is dependent on the *atm-1* apical kinase to induce histone misprocessing. Histone misprocessing triggered by *rpoa-2* depletion is also partially dependent on *gen-1*, and LIN-35 that is independent of *cdl-1* expression. There are likely additional unidentified regulators (?) that remain to be identified. Some illustrations shown were obtained from the NIAID NIH BioArt Source.

## Discussion

Histone proteins are structural components of the chromatin that controls the organization and accessibility of DNA, which are critical for global control of gene expression and developmental homeostasis (55). Several studies in human cells have shown that defects in histone pre-mRNA 3’ end processing can be triggered by cell cycle arrest or after DNA damage (56, 57). Recently, evidence of histone post-transcriptional 3’ misprocessing and subsequent aberrant polyadenylation has also been demonstrated in various pathological states including in cancer cells and neurodegenerative models (18, 19, 58). These evidence provides support that histone 3’ misprocessing is associated with aberrant cell cycle progression; however, we currently do not know if histones are misprocessed as a consequence or a driving factor for pathogenesis. Here, we provide a direct link between rRNA biogenesis and histone homeostasis in *C. elegans* by identifying that genetic disruption to *rpoa-2* can trigger histone 3’ misprocessing in a *cdl-1* dependent and independent manner that engages the conserved genome surveillance pathways (**Fig. 7**).

### Interaction of rRNA biogenesis and CDL-1 in development

In *C. elegans*, rRNA biogenesis inhibition is best characterized by its antagonistic pleiotropic effect where *rpoa-2* depletion in adulthood leads to lifespan extension, but inhibition at the larval stage causes developmental arrest (22, 23). While deficiency in ribosome production driven by the decline in rRNA transcription is a main contributor to growth arrest in *rpoa-2* deficient animals, our data suggest that the broad down-regulation of cell cycle regulating genes likely also contributes to developmental defects. This is supported by our finding that *rpoa-2* depleted worms fail to develop the normal 30-34 intestinal nuclei through endomitosis (59). Furthermore, the requirement for *rpoa-2* in cell division is more clearly demonstrated in the *C. elegans* germline where depletion of *rpoa-2* prevented meiotic entry and contributed to sterility by inhibiting germ cell progression to oogenesis (24). Given that intestinal endomitosis occurs within the first 12 hours at the L1 stage (59), we suspect that the partial binucleation defect we observed is likely due to a delay in RNAi feeding effectiveness in *C. elegans,* where maximal penetrance is achieved typically after at least 24 hours of exposure (60).

We show that while knockdown of *rpoa-2* decreases CDL-1 protein expression, there is also a significant decrease in *cdl-1* gene expression at the mRNA level. This suggests that the effect of *rpoa-2* knockdown on CDL-1 dependent histone misprocessing is not solely caused by a decrease in protein translation capacity from reduced polysome assembly previously reported in *rpoa-2* depleted worms (22). Recently, it was also shown that *rpoa-2* depletion directly alters chromatin accessibility independently of ribosome assembly to influence developmental progression (24). However, analysis of the ATAC-seq data from that study did not show significant changes to the promoter accessibility of *cdl-1* after *rpoa-2* depletion; as such, the precise mechanism of how *rpoa-2* supports *cdl-1* expression at the mRNA level remains to be determined.

At the transcriptome level, our data place *rpoa-2,* at least in part, upstream of *cdl-1* to regulate cell cycle genes; however, we also observed that they co-regulate a subset of genes that was revealed only when both factors were co-depleted. This synthetic interaction on the transcriptome revealed functional redundancy between *rpoa-2* and CDL-1 in regulating a broad range of genes involved in RNA metabolism and nuclear or mitochondrial transport. This highlights a complex genetic interaction between *rpoa-2* and CDL-1 in regulating gene expression in a linear manner, but also in a functional buffering capacity to coordinate the expression of a different subset of genes.

### DNA damage signaling and histone processing defects

A hallmark of RNA Pol I disruption is the onset of nucleolar stress which can manifest as structural changes to the nucleoli and impairment of ribosome biogenesis (61). The canonical response to nucleolar stress is the stabilization of the p53 protein by releasing it from MDM2 inhibition to trigger cell cycle arrest (62, 63). Despite the lack of MDM2 protein in *C. elegans*, disruption of nucleolar proteins has been shown to activate p53/CEP-1 to enhance innate immunity against bacterial pathogens, suggesting some conservation of the p53 dependent nucleolar stress response (64). However, our data showed that histone misprocessing triggered by *rpoa-2* depletion was not dependent on *cep-1*; this suggests that nucleolar stress induced p53 signaling in cell cycle regulation is not shared in worms, which may be consistent with the canonical mechanism signaling through the MDM2 protein that is not conserved in *C. elegans*.

Revisiting upstream regulators of genome surveillance, we found that *rpoa-2* knockdown induced histone misprocessing and *cdl-1* down-regulation through the apical DNA damage response kinase *atm-1*. While ATM-1 activation has not been widely reported to respond to nucleolar stress, a previous study has shown that hypoosmotic stress induced DNA damage in the nucleoli can activate an ATM-1 dependent DNA damage response (65). Consistently, a recent study has also shown that *rpoa-2* depletion activates ATM-1 in the *C. elegans* germline as evidenced by the increase in phosphorylation of the ATM/ATR substrate (24). To our surprise, while *atm-1* mutation completely restored *cdl-1* expression upon *rpoa-2* knockdown, *his-16* encoding the H2A variant was still misprocessed, indicating that *rpoa-2* also regulates histone processing in a *cdl-1* independent manner.

Investigating additional cell cycle regulators revealed that *rpoa-2* dependent histone misprocessing also signals through the Holliday junction resolvase homologue *gen-1* and the Rb homologue protein LIN-35 (54, 66). The *C. elegans gen-1* has been shown to promote cell cycle arrest signaling upon DNA damage, and the LIN-35 protein functions as the pocket protein within the DREAM complex that exerts repression of cell cycle genes (52, 67). Interestingly, *gen-1* and LIN-35 were selectively required for misprocessing of different histone transcripts, illustrating that disruption of *rpoa-2* triggers histone misprocessing via multiple downstream effectors. Perhaps the most surprising finding was that loss of *gen-1* and LIN-35 can suppress histone misprocessing after *rpoa-2* knockdown despite reduced *cdl-1* expression. Previous studies in human cells have shown that histone misprocessing can occur by impaired recruitment of the 3’ processing factors to the stem loop rather than direct SLBP down-regulation (68). Whether *gen-1* and LIN-35 are involved in the recruitment of histone processing factors remains to be determined. Nonetheless, our findings that *rpoa-2* depletion signals through *atm-1*, *gen-1*, and LIN-35 to induce histone misprocessing in a *cdl-1* dependent and independent manner suggest that multiple genome surveillance pathways converge to promote an adaptive state that restricts cellular growth and proliferation in response to the onset of rRNA or nucleolar stress. The engagement of *cep-1* independent factors also suggests that *C. elegans* relies on an alternate nucleolar stress signaling pathway to affect cell growth and development that is non-canonical to the p53 pathway observed in mammals (69).

### Integrator complex in histone 3’ processing

One surprising aspect of our RNAi screen was that we did not identify any novel 3’ cleavage factors, and a subsequent rescreening to a RNAi sub-library of known *C. elegans* cleavage factors genes including *cpsf-3* that cleaves the human histone mRNA also did not yield any positive hits (data not shown). We did however identify *ints-4* encoding part of the catalytic core of the Integrator complex as one of the strongest regulators of histone 3’ processing. RNAi knockdown of *ints-4* caused histone misprocessing to a degree that was greater than that of *rpoa-2* or *cdl-1* depletion. We and others have previously reported that the Integrator complex functions to cleave snRNA in a post-transcriptional mechanism that is analogous to histone 3’ processing in *C. elegans* (5, 70, 71). In human cells, it has also been demonstrated that the Integrator complex contributes to RNA Pol II transcription termination to regulate expression of protein coding genes (72). Knockdown of the Integrator complex or its associated proteins has previously been shown to cause misprocessing of histone transcripts in human cells; mechanistically, it has been proposed that this effect is due to the loss of the histone locus body and the U7 snRNA upon INTS4 depletion (39). Curiously, *C. elegans* lack both the histone locus body and U7 snRNA, which implies that the Integrator may contribute to histone 3’ processing via an independent mechanism. One possibility is that depletion of *ints-4* causes a defect in transcriptional termination that manifests in histone transcript read-through error and cleavage failure, though Integrator functioning as a termination module has not yet been demonstrated in *C. elegans.* Alternatively, given INTS-4 that resembles a Symplekin like protein which serves as a physical scaffold to organize and anchor cleavage factors (73), it may also contribute to histone processing by serving as an essential binding partner for different proteins that may exhibit endonuclease activity towards the 3’ end of the *C. elegans* histone transcripts.

### Limitation and conclusions

A caveat to our study is that we have not established a direct relationship between histone misprocessing and cell cycle arrest beyond demonstrating that intestinal cells undergo endomitosis defects and aberrant expression of cell cycle markers. Additionally, it remains unclear why histone variants employ different regulators for 3’ processing. One possibility with respect to the H3 variant is the divergent putative HDE motif, which may involve the recruitment of unique recognition factors. Nonetheless, our work has highlighted the requirement of rRNA biogenesis in histone 3’ pre-mRNA regulation in *C. elegans*, and describes a mechanism where depletion of the core subunit of RNA Pol I relays a signal through the conserved genome surveillance regulators that is independent of p53/*cep-1* to trigger histone misprocessing. This alternative signaling mechanism may contribute to our understanding of how nucleolar stress can induce cell cycle disruption in the absence of p53, as demonstrated by cell death induced by Dactinomycin treatment in neuroblastoma and lymphocytic leukemia cells that have mutated or deleted p53 (74, 75). Overall, this study uncovered new information on genetic factors that contribute to histone 3’ processing homeostasis in *C. elegans* and implicated conserved genome surveillance regulators in response to nucleolar stress signaling.

## Acknowledgements

Some strains were provided by the *Caenorhabditis* Genetic Centre (University of Minnesota, Minneapolis, MN) which is supported by the NIH Office of Research Infrastructure Program (P40 OD010440). We thank Dr. Guangshuo Ou (Tsinghua University) for sharing the GOU3941 worm strain.

## Data Availability

All dataset supporting this manuscript is found shown in **Table S3**. RNA-sequencing data generated from this study are available on the NCBI GEO repository GSE329687: https://www.ncbi.nlm.nih.gov/geo/query/acc.cgi?acc=GSE329687. Formatted RNA-seq data are shown in **Table S2**.

## Contributor Role Taxonomy (CRediT)

**MGN**: Conceptualization, Formal analysis, Investigation, Methodology, Writing – original draft. **SRD**: Investigation, Methodology, **ACM**: Investigation, Methodology. **CWW**: Conceptualization, Formal analysis, Funding acquisition, Investigation, Methodology, Supervision, Writing – original draft.

